# Molecular architecture of human dermal sleeping nociceptors

**DOI:** 10.1101/2024.12.20.629638

**Authors:** Jannis Körner, Derek Howard, Hans Jürgen Solinski, Marisol Mancilla Moreno, Natja Haag, Andrea Fiebig, Shamsuddin A. Bhuiyan, Idil Toklucu, Raya Bott, Ishwarya Sankaranarayanan, Diana Tavares-Ferreira, Stephanie Shiers, Nikhil N. Inturi, Anna Maxion, Lisa Ernst, Lorenzo Bonaguro, Jonas Schulte-Schrepping, Marc D. Beyer, Thomas Stiehl, William Renthal, Ingo Kurth, Theodore Price, Martin Schmelz, Barbara Namer, Shreejoy Tripathy, Angelika Lampert

## Abstract

Human dermal sleeping nociceptors display ongoing activity in neuropathic pain, affecting 10% of the population. Despite advances in rodents, a molecular marker for these mechano-insensitive C-fibers (CMis) in human skin remains elusive, preventing targeted therapy. In this translational Patch-seq study, we combine single-cell transcriptomics following electrophysiological characterization with single-nucleus and spatial transcriptomics from pigs and humans. We functionally identified CMis in pig sensory neurons with patch-clamp using adapted protocols from human microneurography. We identified oncostatin-M-receptor (OSMR) and somatostatin (SST) as marker genes for CMis. Following dermal injection in healthy human volunteers, oncostatin-M, the ligand of OSMR, exclusively modulates CMis. We identified the entire molecular architecture of human dermal sleeping nociceptors, providing new therapeutic targets and the basis for a mechanistic understanding of neuropathic pain.

**One Sentence Summary:** We identify the molecular architecture and specifically OSMR and SST as molecular markers for human dermal sleeping nociceptors, key players in the generation of neuropathic pain.

**Short version:** In this Patch-seq study, we identify OSMR and SST as molecular markers for human dermal sleeping nociceptors, key players in the generation of neuropathic pain.

## INTRODUCTION

Neuropathic pain, which affects 10% of the population and 20-30% of diabetic patients, correlates with increased activity of sensory neurons in the skin called by the fairly misleading name “sleeping” or “silent” nociceptors (*1–5*). They are defined in human microneurography and psychophysics studies of the skin as mechano-insensitive C-fibers (CMi-fibers (*6*), also termed type 1b fibers (*7*)), and show pronounced chemical responsiveness, e.g. to capsaicin, ATP, partly to histamine, and several inflammatory mediators (*8–11*). Under inflammatory and other painful conditions, CMi-fibers can be sensitized to mechanical stimuli, becoming similar to polymodal nociceptors (CM-fibers or type 1a-fibers (*12*)).

Human skin contains CMis that can be reliably identified by their distinct functional and biophysical properties (*13–16*). In contrast, rodents lack these homologous neurons in their skin; instead, similar neurons primarily innervate internal organs (*17*). The few mechano-insensitive C-fibers identified in rodent skin display significantly different biophysical properties compared to human CMis (*18, 19*). While rodent models crucially advance our understanding, they limit our ability to deduce molecular markers or new drug targets relevant for human skin sleeping nociceptors (*20*). In contrast, pig skin is innervated by neurons resembling human CMis showing tight correlations among mechanical, chemical and biophysical properties (*21, 22*). This similarity provides a valuable translational model for studying dermal nociceptive fibers relevant for human neuropathic pain. In mice, mechano-sensitivity appears less tightly correlated to biophysical properties than in human and pigs, and the functional roles of chemosensing and disease-related hyperactivity may be assumed by other nociceptor classes (*23*).

CMi-fibers are insensitive to strong mechanical stimulation unless sensitized and are unable to follow high-frequency electrical pulses, even when sensitized (*16*). These fibers preferentially respond to slow depolarization, such as low frequency sinusoidal pulses rather than square pulses (*13*). A characteristic feature of these fibers is their enhanced activity-dependent slowing (ADS) upon repetitive stimulation: when stimulated electrically with 2 Hz pulses in the receptive field for an extended time, the conduction velocity of action potentials (APs) traveling along the nerve fiber is reduced (*22*). The distinct electrophysiological characteristics of CMis likely stem from their unique molecular architecture (*24*), potentially offering an opportunity to identify specific, conserved markers for human CMis. In this study, we characterize the molecular architecture of these nociceptors in pigs and humans identifying oncostatin-M receptor (OSMR) and somatostatin (SST) as marker genes for CMis.

## RESULTS

### An integrated multi-modal taxonomy for pig peripheral sensory neurons

Single cell transcriptomic studies of peripheral sensory neurons have revealed a molecular taxonomy of highly specialized and diverse cells responsible for various aspects of sensory perception like pain (*25–30*). In contrast, functional single cell in vitro patch-clamp studies have not yielded a similarly comprehensive cellular taxonomy based on sensory neuron excitability, especially of large mammals given their described difference in electrophysiology compared to rodents (*31*). Recent Patch-seq techniques, however, provide a unique opportunity for bridging the molecular identities of dorsal root ganglion (DRG) neurons with their electrophysiological and morphological identity (*32–37*): individual neurons are electrically characterized via patch-clamp, followed by single cell RNA sequencing.

We used Patch-seq to characterize electrophysiological and transcriptomic characteristics of pig DRG neurons, intentionally targeting smaller, likely C-fiber neurons. We assessed up to 27 electrophysiological features per cell and visualized cellular morphologies prior to capturing whole-cell mRNA (Fig. S1). We successfully sampled 288 neurons using our improved harvesting protocol (Fig. S2, methods) and analyzed their transcriptome by Smart-seq2 and deep sequencing. Of these, 226 neurons passed rigorous quality control of transcriptomes, with a median of 700,000 reads and 11,000 unique detected genes per cell (Fig. S3), demonstrating the deep transcriptomic characterization provided by this methodology.

We next applied droplet-based snRNAseq using the 10XGenomics Chromium v3 platform to sample transcriptomes from 16,997 nuclei from flash-frozen pig DRGs. Following rigorous transcriptomic filtering, we obtained 2,176 DRG neuronal nuclei, with a median of 4,574 detected Unique Molecular Identifiers (UMIs) and 2,131 genes expressed per nucleus. Following unsupervised clustering of snRNAseq-based transcriptomes we identified 16 DRG neuronal subtypes based on distinct marker gene expression (Fig. 1A, see methods). Important pain-related genes, such as *CALCB*, or the receptor for oncostatin M (*OSMR)* were reliably quantified, while a few, including *SCN10A* and *TRPV1*, suffered from some degree of genome annotation incompleteness in the pig, as described previously (Fig. S4, (*38*)). Despite this, TRPV1 expression was detected in expected neuronal subtypes (Fig. S5), ensuring that these genome annotation limitations did not impede our ability to identify its presence where anticipated.

**Figure 1:**
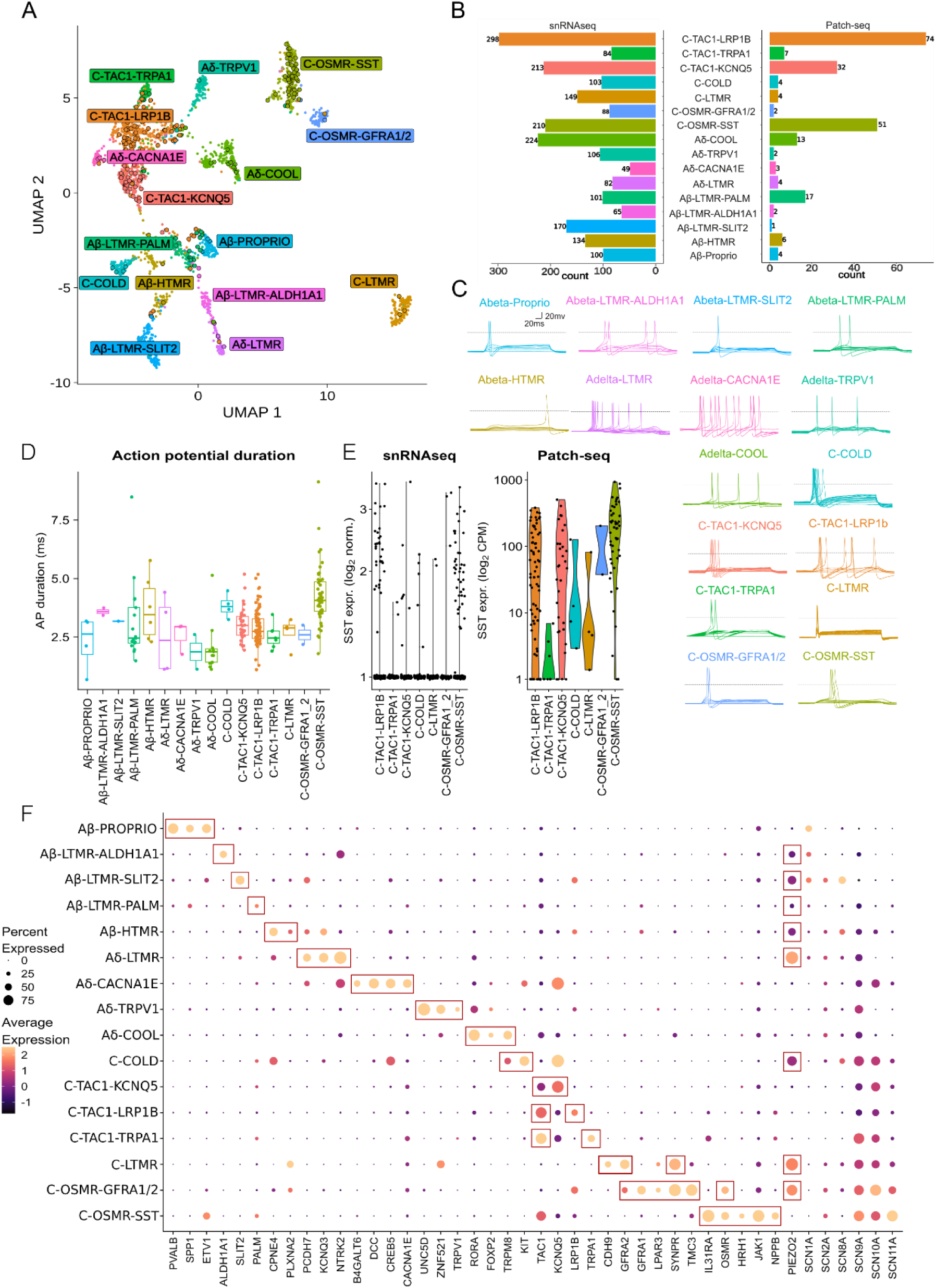
A multi-modal taxonomy of pig dorsal root ganglion neurons. (**A**) Multi-modal neuronal taxonomy of porcine DRG, displaying DRG neuronal subtypes categorized by multi- variate gene expression profiles from snRNAseq (small dots) and Patch-seq (larger circles). Colours indicate neuronal identity based on unbiased clustering of snRNAseq data. (**B**) Overall counts of cells in each neuronal subtype from snRNAseq (left) and Patch-seq (right). **C**) Representative current-clamp voltage traces for exemplar cells characterized via Patch-seq for each cell type in A). Voltage trace shows an overlay of 200 ms rectangular depolarizing pulses of increasing intensities. Scale bar insets represent 20 ms (horizontal) and 20 mV (vertical). Dashed line indicates 0 mv. (**D**) Distributions of action potential duration (C-fibers: 3.34 ± 0.082 ms, A-fibers: 2.75 ± 0.196 ms, *p*=8.88×10−5, two-sided Wilcoxon rank-sum test. Longest AP duration for C-OSMR-SST: 4.26 ± 0.170 ms, p = 6.69 ×10^-11^, two-sided Kruskal-Wallis test for each neuronal subtype (mean ± SEM). (**E**) SST expression in snRNAseq (left) and Patch-seq (right). Each point represents a single cell, with expression quantified by RNA-seq read counts and displayed on a log-transformed y-axis for snRNAseq (left) and Patch-seq (right). Difference in units between axes related to technology differences. (**F**) Group dot plot illustrating snRNAseq-based expression of specific genes for each neuronal subtype. Inset boxes reflect representative markers or cell type-relevant genes discussed in the text.

We mapped each Patch-seq characterized cell to our larger snRNAseq atlas by gene expression guided integration of the two datasets (Fig. 1A, see methods). Each transcriptomically defined sensory neuron subtype was represented by at least one and up to 74 Patch-seq characterized neurons (Fig. 1B). The integration of snRNAseq and Patch-seq datasets effectively mitigated technical differences between the two approaches, as evidenced by the uniform distribution of cells from different batches and biological replicates across the integrated UMAP space (Fig. S6). While we observed some expected differences in injury-induced and culturing-related genes between the datasets (*39, 40*) (Fig. S7), cell-type enriched marker genes were largely consistent between either technology (Fig. S8). Patch-seq cells which are mapped to the same molecular cell type display similar electrophysiological properties (Fig. S1). To corroborate our findings, we also performed “spot-based” spatial transcriptomics (Visium technique) from pig DRG neurons (Fig. S9), using the same approach as reported before for human DRG neurons (*29*), with identification of 5,577 putative neurons with a median of 4,597 detected UMIs and 2,138 expressed genes per spot (Fig. S10). Our thorough taxonomy reflects all major expected neuronal DRG populations, consisting of five Aβ-, four Aδ-, and seven C-fiber related neuronal subtypes (Fig. 1, supplement table 1). In naming the cell types in this study, we designated putative fiber types based on gene expression patterns, knowledge on fiber function in large mammals and through comparisons with existing transcriptomic datasets from large mammals, such as humans and monkeys (*26, 27, 29*). With our approach, we were able to map distinct electrophysiological and morphological features to the transcriptomic identities of the sensory neurons (Fig. 1C, Fig. S1 and S9), e.g. the groups related to C-fibers compared to A-fibers exhibited significantly longer action potential (AP) durations (Fig. 1D) and as expected, had smaller mean diameter cell bodies (Fig. S9).

The Aβ-group of putative proprioceptors, Aβ-Proprio, displayed high expression of *PVALB*, *SPP1* and *ETV1*, hyperpolarized after-hyperpolarization (AHP) minimum (-81.6 ± 1.22 mV) and hyperpolarized AP thresholds (-55.2 ± 4.31 mV) and large cell diameters (95 ± 3.1 µm) (Fig. 1F and Fig. S1, S9). We identified three low-threshold mechanoreceptor subtypes (Aβ-LTMRs), termed Aβ-LTMR-ALDH1A1, Aβ-LTMR-SLIT2, and Aβ-LTMR-PALM, each expressing *PIEZO2* and showing low expression of the pain related sodium channel genes *SCN9A*, *SCN10A* and *SCN11A*. The Aβ-fiber neuron group named Aβ-HTMRs likely represent high-threshold mechanoreceptors, marked by *CPNE4* and *PLXNA2*, as well as *PIEZO2*, and *SCN10A*, suggestive of a potential nociceptive function (*29*).

We identified four Aδ groups: the Aδ-LTMR, with uniquely high expression of *PIEZO2* and marked by *PCDH7*, *KCNQ3*, *NTRK2*; the Aδ-CACNA1E, marked by *B4GALT6*, *DCC*, *CREB5*, *CACNA1E*, and - unique among A-fibers - by its relatively high expression of *SCN10A*; the Aδ- TRPV1, marked by *UNC5D* and *ZNF521*, and with distinctively high expression of the capsaicin and heat receptor *TRPV1*. The Aδ-COOL group was marked by the cold and menthol sensitive ion channel *TRPM8*, *RORA* and *FOXP2* expression. They displayed a very short AP duration (2.01 ± 0.286 ms, Fig. 1D), low AP amplitudes (97.5 ± 2.31 mV) and the highest fraction among all DRG cell types for ongoing activity (61.5%, Fig. S1), potentially due to patch clamp being performed at room temperature which represents a tonic cold stimulus for those cells.

The C-fiber related groups all displayed high expression of the pain-related sodium channel genes *SCN9A*, *SCN10A* and *SCN11A*. C-COLD, was marked by *TRPM8* and *KIT* and showed moderate *PIEZO2* expression, suggesting potential mechano-sensitivity. We identified three C-fiber related subtypes with high expression of *TAC1*, encoding for the nociception-related neurotransmitter substance P, and each subtype showed APs with large amplitudes (Fig. S1).

These three populations differed by strong expression of the potassium channel *KCNQ5* (C- TAC1-KCNQ5), the low-density lipoprotein receptor-related protein 1B (*LRP1B*; C-TAC1- LRP1B), and the pain relevant transient receptor potential channel *TRPA1* (C-TAC1-TRPA1). One C-fiber subtype was marked by *CDH9* and displayed high expression of *PIEZO2* and *GFRA2*; we termed these cells C-LTMRs, given expression of similar markers in a putative orthologous DRG cell type described in humans and mice previously (*29, 30*). We identified an additional group of C-fibers, termed C-OSMR-GFRA1/2, expressing the receptor for oncostatin-M (OSMR) and marked by expression of *GFRA1*, *GFRA2*, *SYNPR*, and *TMC3*, and with high *PIEZO2* expression, suggestive of potential mechano-sensitivity. Finally, we identified a distinct C-fiber subgroup, notable for its high expression of *OSMR*, low expression of *PIEZO2*, and marked by *IL31RA*, *HRH1* (the histamine receptor), *JAK1*, and *NPPB*. We named this subgroup C-OSMR-SST, given the prominent expression of *SST* among these cells, especially among our whole-cell datasets of these neurons (i.e., Patch-seq and spatial transcriptomics, Fig. 1E, Fig. S11). Notably, these cells expressed the highest levels of *SCN11A* and displayed the longest AP duration of any DRG cell type in our dataset (Fig. 1D).

In summary, our pig DRG neuronal taxonomy displays marked neuronal diversity in transcriptomic and electrophysiological characteristics.

### Comparison of pig DRG transcriptomes to cross-species atlases (including human)

Transcriptomic studies of sensory neurons have shown evolutionary conservation of major cell identities between species (*25, 41*). Consequently, to contextualize our pig DRG cell type taxonomy in relation to homologous cell types defined in other species, including humans, we compared the transcriptional identities from our pig snRNAseq dataset with a recently published cross-species atlas of the mammalian DRG (Fig. 2A)(*41*). Following the projection of our pig snRNAseq dataset onto this atlas (see Methods), we observed a good representation of pig neuronal nuclei among the cell type clusters defined in this broader cross-species atlas (Fig. 2B, 2C).

**Figure 2:**
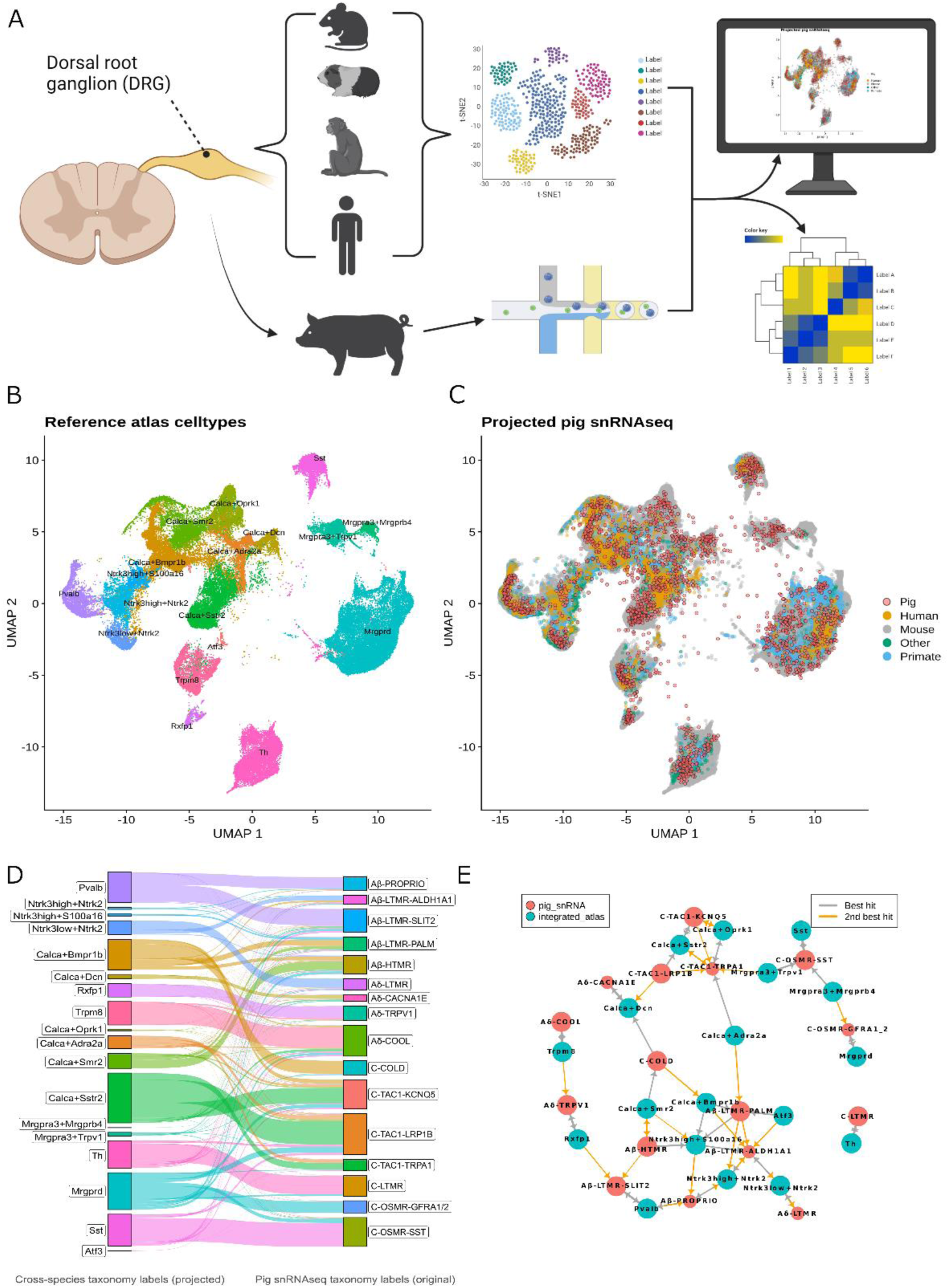
High concordance between pig and cross species neuronal taxonomies. **(A**) Schematic overview of the workflow to compare pig and cross-species neuronal taxonomies. (**B**) UMAP visualization of cell types in the cross-species DRG atlas, with each cluster (colors) representing a distinct DRG neuron cell type. (**C**) Projection of pig snRNA-seq cells (red dots) onto the cross-species atlas (same as A, cells colored by species of dataset origin). (**D**) River plot illustrating the cell type correspondence of pig snRNAseq cells after projecting into cross-species taxonomy (left) and original pig snRNA-seq taxonomy labels (right, cell type colors same as in Figure 1). Width of connecting line strips indicates the count of pig snRNAseq cells mapping between categories. (**E**) MetaNeighbor cluster graph visualization showing the relationship between pig snRNA-seq cell types (red nodes) and cross-species taxonomy cell types (blue nodes). Edges represent strong similarities between cell types, with thicker edges indicating stronger associations. Best hits (grey lines) indicate closest cell types by co-expression whereas 2^nd^ best hit (orange lines) indicates second closest cell type.

To quantify the comparability between the pig and cross-species DRG taxonomies, we employed two complementary analytical approaches. First, we used a Seurat-based label transfer method to project the cross-species atlas cell type labels onto our pig snRNA-seq data. This approach predicts cell type labels for query cells based on transcriptional similarities to the reference dataset. We visualized these results using a river plot (Fig. 2D), which illustrates the flow of cell type assignments from the cross-species classification to our original pig taxonomy. Second, we applied the MetaNeighbor algorithm (39) to evaluate the replicability of cell types across taxonomies. This method quantifies the similarity of cell types based on shared gene expression patterns. We represented these MetaNeighbor scores using a clustergraph plot (Fig. 2E), where edge weights indicate the strength of cell type similarity across taxonomies. For each pig cell type, the top corresponding cell type in the cross-species atlas and its MetaNeighbor area under the curve (AUC) score are provided in Supplementary Table 1.

Overall, we observed excellent concordance between cell type nomenclatures, with several examples of strong one-to-one orthologous relationships (delineated in Supplement table 1), such as between *Sst* (cross-species) and C-OSMR-SST (pig), Mrgprd and C-OSMR-GFRA1/2, Th and C-LTMR, and Adelta-COOL and Trmp8, among others. Additionally, we identified a few one-to-many relationships; for instance, the cell type classified as *Pvalb* within the cross-species atlas corresponded to two cell types in our pig taxonomy, Aβ-PROPRIO and Aβ-LTMR-SLIT2. We also noted that certain cell types, such as Mrgpra3+Mrgprb4 and Mrgpra3+Trpv1, defined in the cross-species atlas largely based on their prevalence in rodents were less frequent in our pig dataset, suggesting that they may be underrepresented in larger mammals like pigs and humans.

While cross-species comparisons offer broad evolutionary insights, they may not fully capture the nuances of larger mammalian systems, particularly for pain-relevant neuronal populations. For instance, the neurons called CMi as characterized by a tight link between lack of mechanosensitivity and specific biophysical properties in human and pig are absent in mouse skin but crucial in human nociception, and *TRPM8*-positive neuron organization differs between rodents and primates (*27*). Given this translational gap to human physiology, we conducted a similar comparative analysis using recent transcriptomic datasets composed exclusively of human DRG cells (*25, 27, 29*), offering a more focused evaluation of the alignment of our pig cell types with those in humans (Fig. S12). Here, we also observed a strong correspondence between the cell types defined in our pig DRG taxonomy and those identified in human-specific atlases.

These analyses further reinforce the notion that the pig serves as a robust model for studying human sensory neurobiology, with pig DRG cell types closely aligning with those found in humans, both within the broader cross-species context and within the more specific framework of human-focused taxonomies.

### Identification of C-OSMR-SSTs as probable sleeping nociceptors

Having described the pig DRG neuronal taxonomy and confirmed its validity to human DRGs, we aimed to identify the subgroup which represents the sleeping nociceptors within those DRG neuronal subtypes. CMi-fibers can be reliably identified in humans and pigs *in vivo* by a combination of three functional electrophysiological characteristics. To transfer this *in vivo* stimulation paradigm to our *in vitro* experiments, we applied our optimized Patch-seq electrophysiological characterization pipeline (Fig. 3).

**Figure 3:**
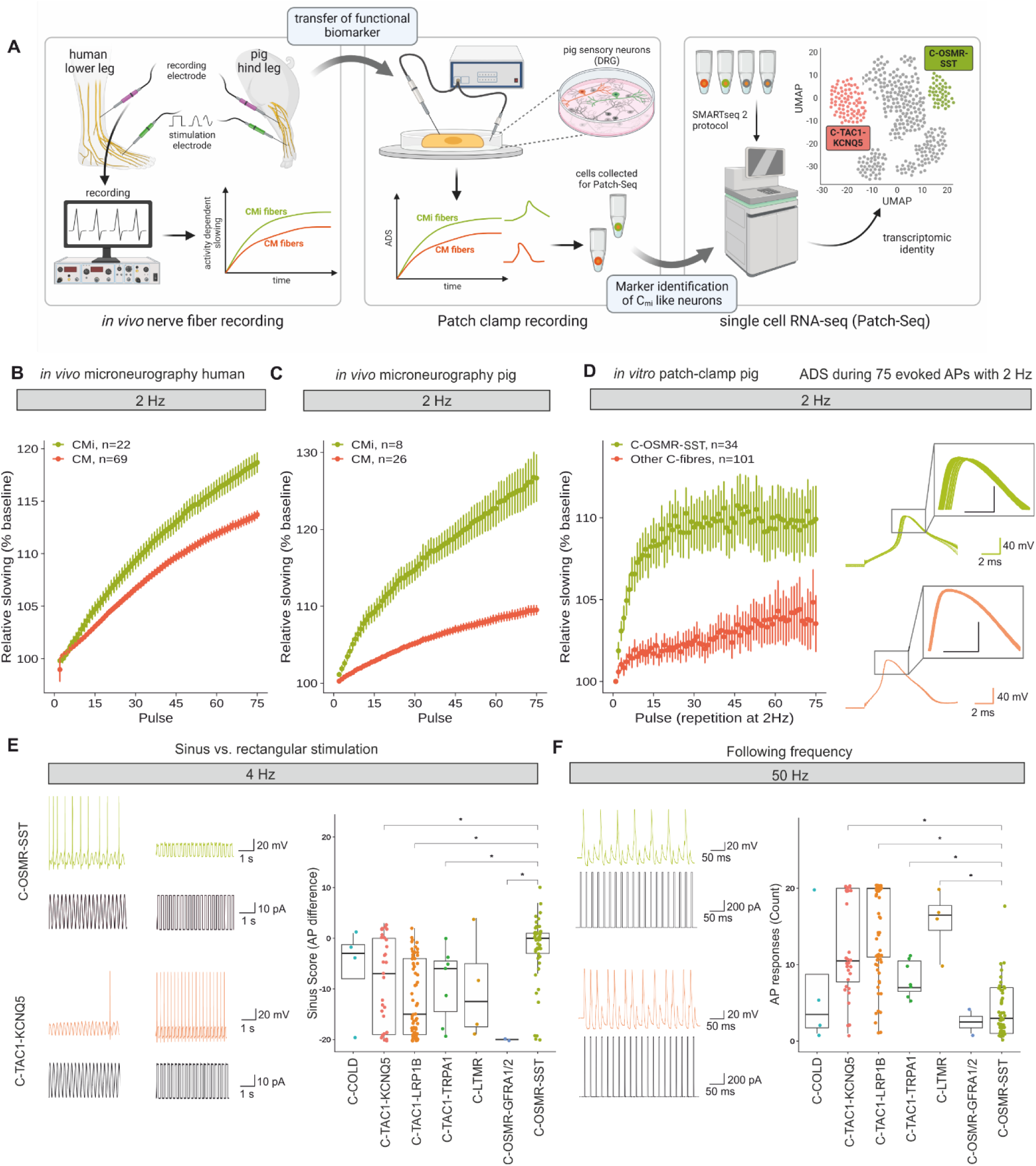
Molecular identity of CMis in Patch-seq and adapted electrophysiological stimulation protocols in vitro. **(A**) Schematic for identifying molecular markers of mechano-insensitive C-fibers (CMis). CMis are traditionally identified *in vivo* using electrical stimulation protocols applied to the skin (left panel). Functional biomarkers of CMis include activity dependent slowing (ADS) upon repetitive stimulation, observed in CMis but not mechanosensitive C-fibers (CMs). We transferred these CMi-discriminating functional biomarkers into electrophysiological stimulation protocols that can be applied to DRG neurons using Patch clamp recordings *in vitro* (middle panel). This enables identifying molecular markers of putative CMi-like neurons using transcriptomics from Patch-seq (right panel). (**B,C**) *In vivo* characterization of ADS (y-axis) relative to stimulation pulse number (x-axis) for CMi-fibers (green) and polymodal CMs (red) in humans (B) and pigs (C). (**D**) Similar to B,C), showing ADS from Patch-seq characterized neurons stimulated using somatic current injection. Cells colored by molecular cell type, highlighting differences between C-OSMR-SST cells (green, 110 ± 2.04%, n=34) compared to all other C- fiber neurons in Patch-seq dataset (red, 104 ± 1.73%, n=110, p = 9.27×10^-07^, Mann-Whitney U = 741.5). Right panel shows overlay of 75 APs induced by 1 nA current injection for 10 ms delivered at 2 Hz from an example C-OSMR- SST (top) and C-TAC1-KCNQ5 neuron (bottom). Note that APs become more delayed over the course of the stimulation protocol, especially in the C-OSMR-SST neurons, but comparatively less so in the C-TAC1-KCNQ5 neurons. (**E**) Left, Representative voltage trace (top) and current stimulation (bottom) traces evoked by sinusoidal (left) versus rectangular stimulation (right). Right, Quantification of neuronal sinus scores across C-fiber neurons characterized via Patch-seq. Sinus scores are defined as the difference in AP count evoked between sinusoidal versus rectangular stimuli; positive scores indicate greater CMi-like activity. ANOVA (F(6, 158) = 9.03, p < 0.00001. (**F**) Left, Illustration of calculation of following frequency, where neurons are stimulated with 20 1nA current pulses for 4 ms at 50 Hz. Right, count of AP responses evoked by 50 Hz stimuli designed to assess efficacy to follow high frequency stimuli; fewer evoked APs indicate greater CMi-like activity. Asterisks denote statistical significance with p<0.05 as determined by an ANOVA (F(6, 166) = 24.59, p < 0.0001) followed by Tukey’s post- hoc HSD test.

*First*, we applied a train of 75 suprathreshold stimulations delivered at 2 Hz to the patched cells to assess activity-dependent slowing (ADS) (n=176 cells) (*42*).High amount of activity, e.g. due to a 2 Hz stimulation in sleeping nociceptors in human or pig skin *in vivo,* induces substantial slowing of conduction velocity (Fig. 3A,B,C, (*22*)). C-OSMR-SST neurons displayed more ADS compared to other C-fiber related classes (Fig. 3D, Fig. S13). During the stimulation, the C- OSMR-SST neurons displayed significant activity-dependent membrane potential hyperpolarization, reduced AP peak and AP maximum upstroke slope relative to other C-fiber cells (Fig. S13A-C).

*Second*, trains of twenty sine wave and square pulses were delivered at 4 Hz (i.e., sinus score, see methods, n=213 cells) to assess the preference for sine waves versus square pulses observed *in vivo* for sleeping nociceptors (*13, 16, 22*). The C-OSMR-SST subgroup demonstrated the highest sinus score (Fig. 3E, *right*).

*Third*, trains of 20 short suprathreshold square pulses were delivered at 50 Hz (n=226 cells, Fig. 3F), as CMi fibers *in vivo* are unable to respond to high frequency stimulation continuously with AP generation (i.e., they have a lower following frequency, (*13, 16, 22*)). The C-OSMR-SST subtype displayed the fewest AP responses, significantly lower than the C-TAC1-KCNQ5, C- TAC1-LRP1B, and C-LTMR subtypes (Fig. 3F, *right*).

To further validate our inference that C-OSMR-SST neurons likely correspond to sleeping nociceptors (CMis), we conducted an independent analysis to determine whether neurons with CMi-like electrophysiological features also express genes overlapping with C-OSMR-SST markers. First, we performed a principal component analysis (PCA) on all included neurons regardless of transcriptomic identity incorporating the three functional hallmarks of CMis— ADS, sine wave affinity, and reduced following frequency at 50 Hz—along with AP duration (Fig. 4A). As expected, the first principal component (CMi ephys PC1) accounted for the majority of variance in these features (49%), indicating that this component could serve as a single score for CMi-like neuronal identity. Leveraging gene expression profiles collected from the same neurons via Patch-seq, we performed a transcriptome-wide analysis to identify genes significantly associated with CMi scores (Fig. 4B). We found a substantial overlap between these electrophysiology-based CMi-associated genes and markers of C-OSMR-SST neurons (Fig. 4C, 76 genes out of 228 total CMi-associated genes, p-value for hypergeometric test = 2.34 × 10⁻⁷⁹). This analysis further highlighted several key pain-related ion channels, including *SCN11A* (Fig. 4D) and *SCN10A*, suggesting that their expression is closely related to CMi-like neuronal characteristics and underscoring the potential future therapeutic value of targeting these channels (see Discussion).

**Figure 4:**
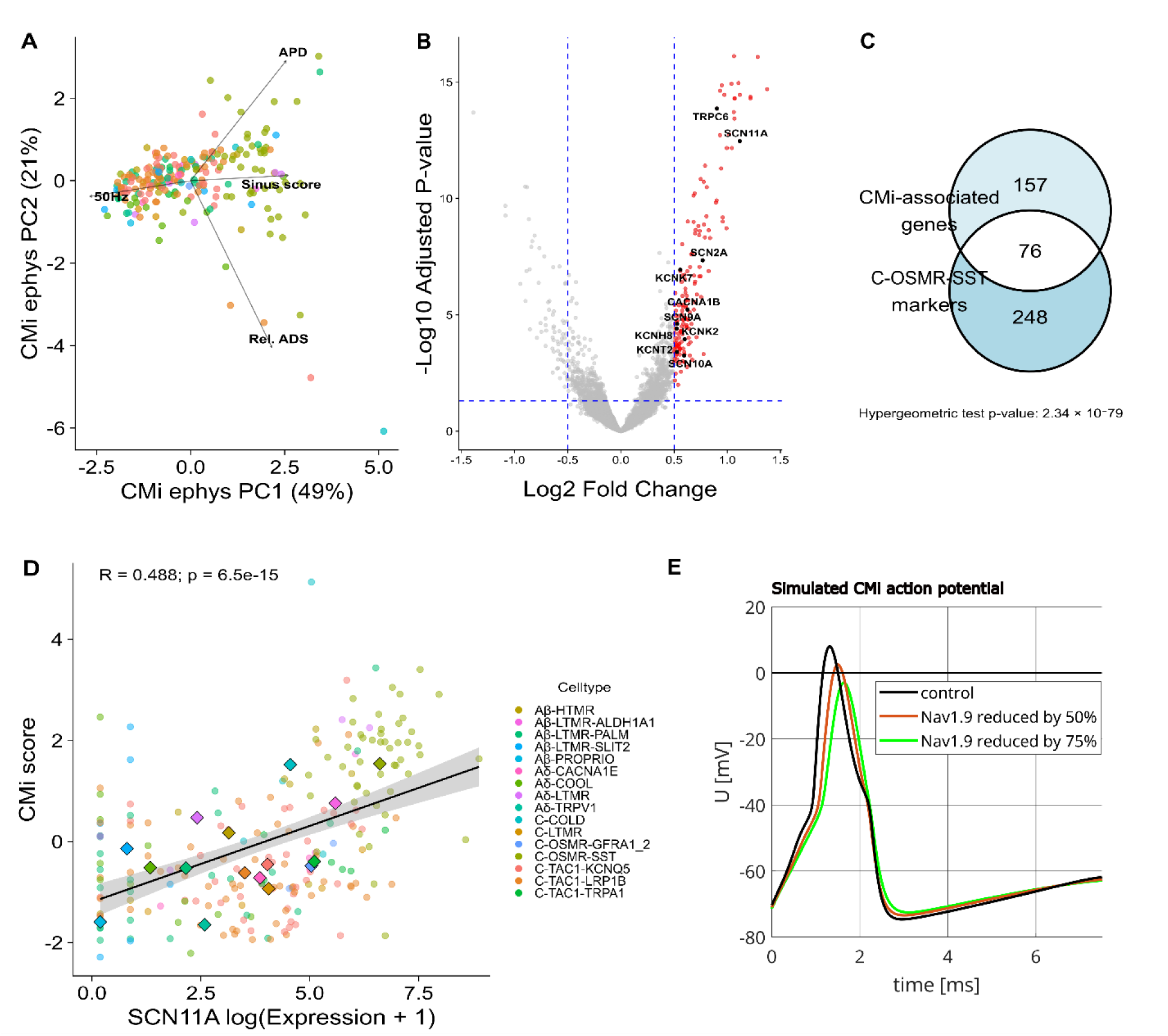
Electrophysiological feature guided analysis of genes associated with CMi phenotype. **(A**) PCA biplot of CMi-related electrophysiology features. Each point represents a single neuron, colored by cell type. Arrows indicate the contribution of each electrophysiological feature to the principal components. (**B**) Volcano plot showing CMi score-associated genes. Axes indicate magnitude (x-axis) and statistical significance (y-axis) of association between gene expression and CMi score. Red dots indicate significantly differentially expressed genes (adjusted p-value < 0.05 and log2 fold change > 0.5). Black dots highlight ion channel genes. Vertical blue dashed lines indicate log2 fold change thresholds, and the horizontal blue dashed line indicates the adjusted p-value threshold. (**C**) Venn diagram illustrating the overlap between CMi-associated genes (red genes highlighted in B) and C-OSMR-SST markers (derived from snRNA-seq data). (**D**) Correlation between SCN11A expression and CMi score. Each point represents a single neuron, colored by cell type. Larger diamond-shaped points indicate the mean expression and CMi score for each cell type. The black line shows linear regression fit with 95% confidence interval. **(E)** Simulated action potential waveforms demonstrating the effect of reducing Nav1.9 (SCN11A) conductance. Black trace shows control condition, while red and green traces show the effects of 50% and 75% reductions in Nav1.9 conductance, respectively, illustrating the channel’s contribution to action potential characteristics.

To establish a mechanistic linkage between the gene expression pattern and the functional phenotype of C-OSMR-SST neurons we used the expression data of these neurons from the snRNAseq dataset to generate a computational model based on an extension of the Hodgkin-Huxley approach by using the modeling framework from (*43*) (see methods). In this model, for instance, reduction of the SCN11A conductance of 50% leads to a strong reduction of the action potential duration from 1.34ms to 1.18 ms (Fig. 4E, Fig. S14). The long action potential duration as well as the action potential slope of C-OSMR-SST neurons was highly dependent on the expression of SCN11A, suggesting this channel is a key player in shaping the characteristic functional phenotype of CMi fibers.

Altogether, our *in vitro* experiments, modeled on adapted *in vivo* microneurography stimulation protocols, demonstrate that C-OSMR-SST neurons exhibit functional properties similar to those of CMi-fibers identified in human skin *in vivo*.

### OSM selectively modifies CMi-fibers *in vivo* in humans

Our experiments suggest that C-OSMR-SST neurons are strong candidates for molecularly defined CMi-fibers in pigs and humans. To show that OSMR is a functional marker selective for human dermal sleeping nociceptors, we applied oncostatin-M (OSM), the activator of OSMR, locally to the skin of healthy volunteers (Fig. 5A). As expected for CMi-activation in human skin *in vivo* (*44*), we observed a widespread axon reflex erythema (i.e., skin redness) 24 h following subcutaneous injection of OSM in four out of four participants (Fig. 5B, 275 ± 108.3 mm^2^).

**Figure 5:**
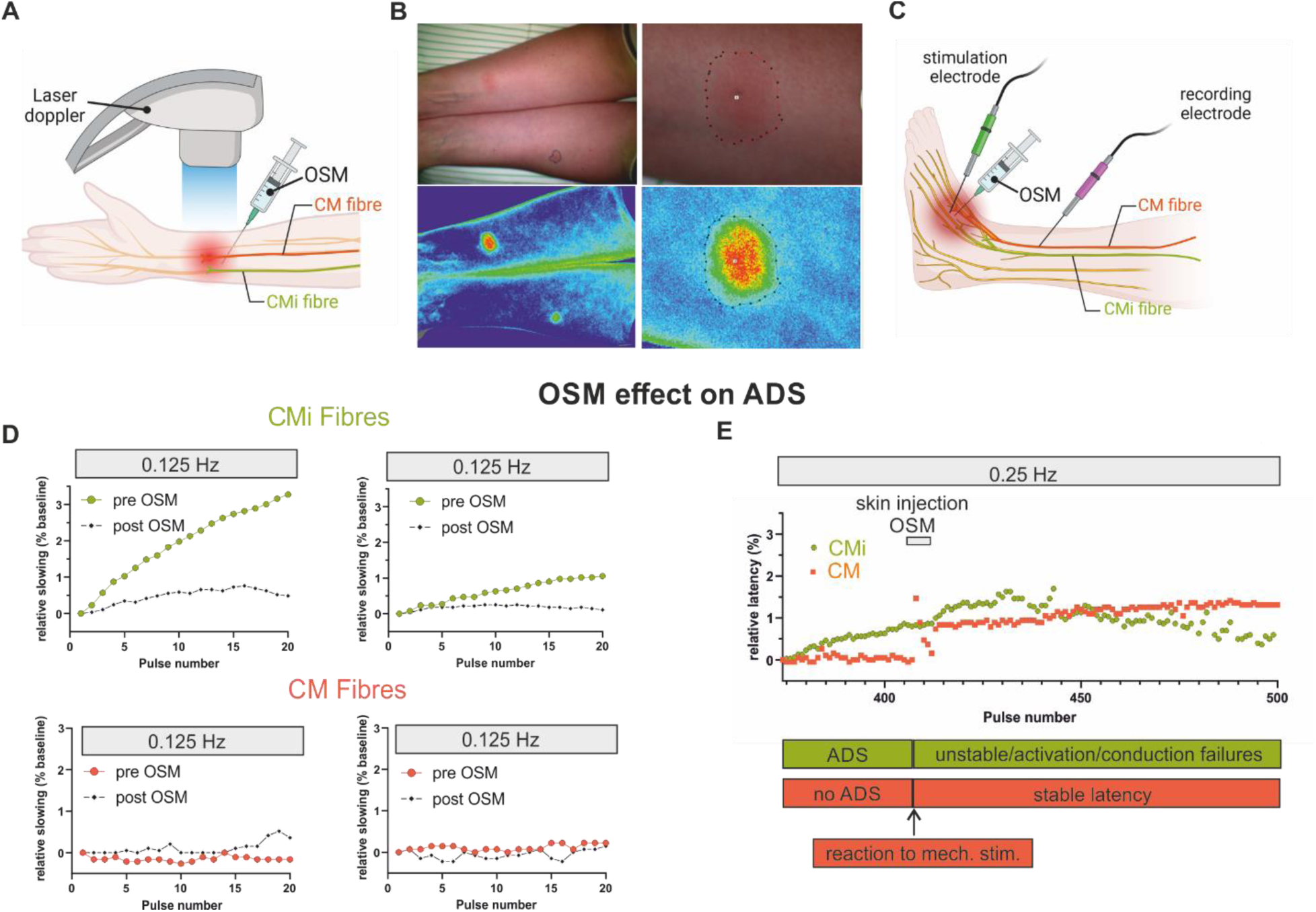
Applying OSM in humans in vivo selectively affects CMi fibers. (**A**) Schematic illustrating the use of oncostatin-M (OSM) during human superficial blood flow measurements. (**B**) Injection of OSM induces an axon reflex erythema of 275 ± 108.3 mm^2^ (n=4) in human skin after 24 h. Images show superficial blood flow (lower) and corresponding photographs (upper) of one participant’s forearms at 24 h after injection (500 ng right distal, 250 ng left proximal). Lower panels show close-ups of injection site in right arm. (**C**) Schematic illustrating the use of oncostatin-M (OSM) during human microneurography experiments. OSM and electrical stimulation are applied to C-fiber nerve endings while AP responses of corresponding C-fiber axon(s) to the constant low frequency electrical stimulation are recorded simultaneously. (**D**) Microneurography recordings of two CMi- and two CM-fibers before (large colored circles) and 7 min after injection of 500 ng OSM (small black markers). Y-axis indicates differences in ADS as change in relative AP latency following delivery of 0.125 Hz test pulses. Following application of OSM, characteristic ADS is abolished in CMis but no change in ADS is observed in CMs. (**E**) Microneurography recordings of a CMi- (green) and CM-fiber (orange) before and after intraepidermal OSM injection during repetitive stimulation with 0.25 Hz. Subsequent response latencies of electrically induced APs are shown normalized to baseline latency determined before pulse 374. The CMi-fiber undergoes ADS during initial pulses. The CM fiber is mechanically activated by injection, as seen by a sudden latency increase followed by normalisation, and afterwards, ADS is stable. In contrast, the mechano-insensitive CMi-fiber is not activated by the mechanical aspect of injection but shows unstable latency afterwards. Some of these instabilities may be occurrence of spurious OSM induced discharges and numerous conduction failures following OSM injection, demonstrating an affection of CMi- but not CM-fibers by OSM injection.

Three participants did not report altered sensation, while one out of four participants experienced moderate itch over several minutes post OSM injection (3 out of 10 on the numeric rating scale). There was no itch or ongoing pain sensation for the next 24 h, apart from mechanical hypersensitivity upon pressure.

We recorded two CMi- and two mechano-sensitive C-fibers (CM-fiber, polymodal nociceptors, type 1a) with microneurography of the superficial fibular nerve in two healthy human volunteers (schematic in Fig. 5C). ADS of two out of two recorded CMi-fibers was strongly reduced, if not abolished, upon intracutaneous injection of OSM in the C-fiber’s receptive field (Fig. 5D). In two out of two mechano-sensitive CM-fibers, on the other hand, OSM injection did not result in any observable ADS change (Fig. 5D).

In one participant, a recorded CMi-fiber shared its receptive field with a CM-fiber. Acutely injecting OSM resulted in an immediate mechanosensory response in the CM- but not the CMi-fiber (Fig. 5E). Seconds after OSM injection, we observed that the latency of the regularly electrically induced APs of the CMi-fiber became unstable and latencies increased, indicating fiber activity (Fig. 5E). Such changes, beyond the initial mechanosensory response, were not observed in the CM-fiber. Based on the response of CMi-fibers – but not CM-fibers – to OSM, we conclude that OSM application results in a selective reduction of ADS in human CMi-fibers but not CM-fibers.

Thus, we provide evidence that CMi-fibers but not CM-fibers in the skin are affected by OSMR application *in vivo*, making OSM-receptor expression by CMi but not CM likely. This suggests that these CMi-fibers are likely to correspond to the molecular subgroup of C-OSMR-SST neurons, identified by our multimodal taxonomy.

## DISCUSSION

The research described here presents a major advance in our understanding of the molecular underpinnings of sensory neurobiology and pain. We have defined a population of neurons consistent with the elusive population of human dermal sleeping nociceptors, i.e., mechano-insensitive C-fibers (CMis, type 1b), by bridging *in vivo* and *in vitro* physiology with multiple single-cell transcriptomic techniques in pigs and humans. The unique strength and unprecedented translational power of our data set lies in the direct link between functional and transcriptomic data for each identified sensory neuron subtype. This allows us to paint the complex picture of the molecular and functional identity of the CMis in human skin, laying the basis for drug target identification, which may counteract their spontaneous activity observed during neuropathic pain but keeps the warning function for potentially tissue damaging stimuli via other nociceptive neurons functional (*1–4*).

C-OSMR-SST, the subgroup identified as CMis, express the highest level of *SCN11A*, coding for the voltage-gated sodium channel Nav1.9, which has been discussed intensely as drug target for neuropathic pain (*45, 46*). We hypothesize that this channel, in combination with other channels expressed in this neuron type, including *SCN10A/*Nav1.8, gives rise to the distinct functional characteristics of C-OSMR-SST neurons, including sine wave sensitivity and formation of very broad APs (*45, 47*). We included the sodium channel expression identified in this manuscript into a basic computer model of CMi-fibers, based on (*43*). We identify the surprising contribution of Nav1.9 to the steepness of the AP upstroke and the shoulder. Our findings thus underscore the prominent role of Nav1.9 in CMi fiber electrogenesis and the promise for developing modulators of Nav1.9 channel function, as these might be especially effective in silencing sleeping nociceptors. It is highly likely that more targets will emerge from our data set, and future studies may show their translational and therapeutic potential.

OSM reduces activity dependent slowing and thus enhances excitability of CMis. Given the involvement of OSM in the JAK pathway (*48*) and the high JAK1 Expression in the C-OSM- SST subgroup, we speculate that the pathway we identified as molecular marker, involving JAK, could be a potentially sensitizing pathway, which may change the CMi-fiber to a more CM-fiber like phenotype during pathophysiological processes. A subset of CMi-fibers, which are histamine sensitive, are associated with itch, and JAK-inhibitors are in clinical use as antipruritic agents (*49*). The specific mechanisms of the contribution of CMi-fibers to itch and neuropathic pain still remains to be elucidated. Interfering with JAK signaling may prevent CMi-fiber sensitization and thus may potentially counteract the development of neuropathic pain and chronic itch beyond its known anti-inflammatory effects (*50, 51*).

Our human experiments using microneurography demonstrate that CMis are responsive to application of OSM, thus substantiating our transcriptomic inference that these cells express OSMR. However, further disentangling how the application of OSM changes CMis, and critically, whether OSM sensitizes CMis in the response to stimuli such as mechanical stimulation, inflammatory agents, cold, heat, remains to be investigated and is beyond the scope of the current study. We recognize that OSM-induced skin erythema in human volunteers stretched out far beyond the actual injection site with blurred borders. Inflammation e.g. after UVB irradiation causes a direct vasodilation independent from neuronal influence, the erythema is – different to our observations - restricted to the site of injury and has sharp borders (*52*), leading to our inference that OSM is directly acting on CMis.

Our analyses demonstrate that the C-OSMR-SST cell type is highly transcriptionally conserved across mammalian species. However, some genes, such as the nicotinic acetylcholine receptor *CHRNA3*, were previously identified as a marker of sleeping nociceptors in mice (*17, 53*), but we did not detect it in either of the three pig datasets presented in this study. In mice this neuronal subgroup is mechano-insensitive in dissociated cultures but expresses nociceptor-related gene products (*17*). These cells have few, if any, cutaneous afferents in mice. In line with this finding, cutaneous sleeping nociceptors have been challenging to detect thus far in murine skin tissues in contrast to humans and pig (*54*). Beside *CHRNA3*, recently, *TMEM100* has been described as a marker involved in unsilencing of *CHRNA3*+ neurons in mice (*53*). As in human sensory neurons (*27*), in our pig dataset *TMEM100* is expressed at very low levels in any neuron type. We thus assume, that the transcriptomic identities of sleeping nociceptors differ between skin and deep somatic tissues, or that CMi-fibers of mice and large, fur lacking mammals such as human and pigs, differ significantly (*17, 54*).

A subset of CMi fibers can be activated by intense heating of the skin, typically at an average surface temperature of 48.0 ± 3.0°C (9). However, this temperature threshold is painful for most individuals, poses a risk of skin damage, and may not generate high enough temperatures in deeper layers of the skin. Notably, intracutaneous injections of capsaicin in human skin activate nearly all CMi fibers (13) suggesting that this fiber class could be triggered by heat stimuli if they could be applied deeply enough in the skin.

As expected with our use of the 10X Genomics Single Cell 3’ Kit, snRNAseq read coverage is biased towards the 3’ ends of gene bodies. Upon qualitative inspection, we observed that some genes, including SCNA10A and TRPV1, exhibit unannotated reads in their 3’ UTRs . We believe this is likely due to incomplete annotation of the 3’ UTRs in the reference transcriptome used.

However, this issue appears to be gene-specific, as other manually assessed genes, such as CALCB, OSMR, and SST, do not show similar evidence of unannotated reads. Despite potential suboptimal annotation for TRPV1 in pigs, we detected TRPV1 transcripts in a significant number of C-OSMR-SST neurons, which aligns with human microneurography data showing heat responsiveness in most CMi fibers.

In designating fiber types such as Aβ-, Aδ-, and C-fibers in our study, we relied on gene expression profiles and comparisons with existing transcriptomic datasets from large mammals as well as on knowledge on fiber function in these species. While this allowed us to align our classifications with well-characterized clusters, we acknowledge that distinguishing certain fiber types, such as Aδ- from C-fibers, remains challenging due to overlapping gene expression patterns and species-specific differences (*55*).

The cell type identified with the highest transcriptomic similarities to the C-OSMR-SST cells are the C-OSMR-GFRA1/2 neurons. The latter, however, show a very negative sine score and no detected ADS arguing against CMi-fiber identity. In our Patch-seq data, we only identified two cells of the C-OSMR-GFRA1/2 population. We note, however, that expression data of this subgroup demonstrates a high *PIEZO2* level in pigs and humans, suggesting mechanosensitivity (*56*). PIEZO2 by itself is not solely responsible for mechanosensitivity in sensory neurons (*57, 58*), but it still indicates that the expressing cells are likely mechano-sensitive CM-fibers.

In conclusion, our work demonstrates the potential of our multi-modal peripheral sensory neuron taxonomy as a powerful tool for identifying molecular markers of cell types that are highly relevant to human pathophysiology, as we showed in the successful molecular identification of human dermal sleeping nociceptors.

## MATERIALS AND METHODS

### Blinding, Power analysis

Besides psychophysical experiments, the study did not contain multiple study groups. Hence, the investigators were not blinded during experiments and outcome assessment. There was no predetermination of sample size with statistical methods. In psychophysical experiments the volunteers were blinded for injection of OSM or control solution.

### Animals

#### Patch-seq dataset and Visium spatial transcriptomics dataset

Dorsal root ganglia (DRG) of pigs were sampled according to the 3R criteria for reductions in animal use, as leftovers from previous independent animal studies (e.g. LANUV reference no. 81-02.04.2018.A051). For this purpose, 10 female pigs of the German Landrace breed, with an average age of 4.5 month and weight of 47.4 kg (SD 5.2 kg) were euthanized either using an overdose of pentobarbital 60 mg/kg body weight or combination of exsanguination in deep anaesthesia and overdose. Subsequently, the DRG were collected as described previously (*59*). DRGs from all segments (cervical/thoracic/lumbar) were included in dataset generation.

#### snRNAseq dataset

Male domestic pigs (German Landrace) were euthanized according to ethical approval obtained from the local authorities (RP Karlsruhe, Germany) under the approval number G-78/18.

Subsequently, thoracic and lumbar DRGs were quickly dissected, washed once in PBS and stored overnight in RNAlater^®^ at 4°C.

| Biological replicate number | Protocol | Modification |
| --- | --- | --- |
| 1 | (26) | None |
| 2 | (57) | + FACS |
| 3 | (57) | + FACS |
| 4 | (58) | None |
| 5 | (58) | None |

### DRG tissue processing

#### Patch-seq DRG preparation

DRG preparation was performed as in (*59*). Briefly, DRG of pigs were transferred on ice and fine excision was performed in ice-cold DMEM F12 medium containing 10% FBS. DRG were treated with 1 mg/ml collagenase P, 1 mg/ml trypsin T1426 and 0,1 mg/ml DNase for digestion.

DRG were cut into small pieces inside the digestion medium for surface enlargement. DRG were incubated in 37°C, 5% CO2 for 120 minutes ± 30 minutes. Approximately after 60 minutes in digestion medium, DRG were triturated using a plastic pipette. After the full incubation time, DRG were triturated three times using glass pipettes with decreasing tip diameter. For further purification, DRG were centrifuged at 500 x G and 4°C twice for four minutes each and the pellets were suspended in DMEM F12 with 10% FBS. DRG were subsequently separated from the lighter cell fragments and myelin by centrifugation of a Percoll gradient containing a 60% Percoll and a 25% Percoll gradient for 20 minutes at 500 x G. DRG neurons were plated on coverslips coated with poly-D-lysine (100 μg/ml), laminin (10 μg/ml) and fibronectin (10 μg/ml). Neurons were then cultured in Neurobasal A medium supplemented with B27, penicillin, streptomycin and L-glutamine and used for voltage-clamp recordings after 12-72 hours in culture.

Addition of recombinant NGF, frequently used in DRG cultures, has been described to decrease the fraction of mechanical insensitive fibers and decrease the amount of ADS in mechanical insensitive fibers (*17, 60*). Therefore, we avoided the use of NGF in all our experiments.

#### snRNAseq DRG preparation

12-16 hours after euthanasia, adjacent fat tissue and the nerve root and spinal nerve were trimmed from the DRGs under a dissecting microscope. The cleaned DRGs were flash-frozen in liquid nitrogen and stored at -80°C until use for nuclei isolation.

Nuclei were isolated from 80 mg thoracic and lumbar DRG tissue per sample employing previously published protocols (*27, 61, 62*) that differ with respect to tissue lysis conditions (Triton X-100 or IGEPAL CA-630) and strategies employed for separation of debris and nuclei (40 µm filtration plus iodixanol density gradient centrifugation or 40 µm filtration plus magnetic-assisted enrichment of neurons using rabbit polyclonal anti-NeuN antibody (Millipore, cat# ABN78, dilution 1:5,000) and anti-rabbit IgG microbeads (Miltenyi Biotech, cat# 130-048- 602, dilution 1:5)). The protocol by Ernst et al. that only relies on 40 µm filtration was modified to include an additional step of fluorescence-assisted enrichment of neuronal nuclei after the final two nuclei washing steps of the protocol (*61*). For this purpose, the nuclei were stained with Anti-NeuN-Alexa Fluor 555 (Millipore, cat# MAB377A5, dilution 1:100) for 30 minutes at 4°C on a rotator and subsequently with DAPI for 5 minutes on ice. At the FlowCore Mannheim, double-positive singlets were sorted into PBS/1% BSA using a BD FACSAria IIu equipped with a 100 µm nozzle. All protocols resulted in similar high-quality nuclei (based on microscopic inspection) and were therefore used as input for microfluidics-based generation of single-nucleus gene expression libraries.

#### Visium spatial transcriptomics DRG preparation

Lumbar DRGs from female domestic pigs (German Landrace) were recovered shortly after euthanasia. The tissue was trimmed of any fat or connective tissue, rinsed once with artificial cerebrospinal fluid, and then freshly frozen by burying it in pulverized dry ice for 1 minute at the time of extraction. All DRGs were subsequently stored in a -80°C freezer. 22 fresh-frozen DRGs from the lumbar region of 7 female pigs were embedded in optimal cutting temperature (OCT) using a cryomold placed over dry ice. To avoid thawing, the OCT was poured in small volumes surrounding the tissue. Two sections of the embedded tissue were cryosectioned at 10 µm, 200 µm apart from each other to ensure different neurons were sliced. The sections were mounted onto SuperFrost Plus charged slides for staining with Eosin and Hematoxylin (HE) or on the capture area of Visium slides for spatial sequencing, while avoiding folding or overlapping.

### Patch-seq Recording Procedures

#### Patch-Solutions

Extracellular solution contained (in mM): 140 NaCl, 3 KCl, 1 MgCl2, 1 CaCl2, 10 HEPES, 20 glucose (pH 7.4; 300-310 mOsm). Intracellular solution contained (in mM): 4 NaCl, 135 K-gluconate, 3 MgCl2, 5 EGTA, 5 HEPES, 2 Na2-ATP, 0.3 Na-GTP (pH 7.25; 290-300 mOsm).

To minimize RNAse contamination of the intracellular solution (ICS), 100x stock solutions of each component (except ATP and GTP) were made using RNAse-free H2O or DEPC-treated and autoclaved where appropriate. Afterwards, the final ICS was manufactured by adding GTP, ATP and RNAse free water. Osmolality control was performed.

#### Patch-Clamp Recordings

Experiments were performed using a HEKA EPC 10USB amplifier and PatchMaster and analyzed using FitMaster v2.8 software (all HEKA electronics, Lambrecht, Germany). Pipette resistance was 1.5-3.5 MOhm. Currents were low-pass filtered at 10 kHz and sampled at 100 kHz. The liquid-junction-potential was corrected for +14.3 mV. All experiments were performed at room temperature. After reaching giga-seal and pipette capacitance correction, the whole cell configuration was established. The resting membrane potential (RMP) was measured immediately after establishing the whole-cell configuration. Holding current was then adjusted to achieve a membrane voltage of -60 ± 3 mV.

The sequence of applied pulse protocols was as follows: First, APs were elicited with 200 ms square pulse depolarisations of increasing intensity. Then, 4 Hz rectangular and 4 Hz sinusoidal depolarization pulses were injected with a 5 second break in between to confirm the threshold and compare also longer rectangular threshold with the sinusoidal threshold (sinus_score). This was followed by 20 4 ms rectangular injections of 1000 pA at 2, 5, 10, 25, 50 and 100 Hz for assessment of the maximum follow frequency. Afterwards, the threshold for 500 ms halfsinus- shaped injections was assessed. Finally, a protocol of 75 1000 pA 10 ms rectangular pulses were injected for assessment of changes in slope, membrane potential, AP peak and rising time changes upon repetitive stimulation as a measurement of axonal slowing (*42*). The electrophysiological assessment of one cell takes around 11 minutes.

#### mRNA harvesting for Patch-seq

Given the large size of pig DRG neurons, the sample collection followed a procedure using two separate pipettes as follows (Fig. S1): First, the recording pipette was retracted under slightly negative pressure from the cell and immutably broken into in PCR-tube containing 4 µl of lysis buffer (40 mM Guanidine hydrochloride, 0.1 mM Smart dT30VN primer, 5 mM dNTPs), secondly a second larger pipette with diameter customized for cell diameter of the cell to collect (resistance ∼0.7 Mohm) was used to collect the entire cell. After collection, this pipette was broken into the same PCR-tube. Samples were immediately frozen on dry ice. Complete and exclusive collection was documented with the microscope’s camera.

#### Extraction of electrophysiological features

Electrophysiological feature extraction was performed using in-house scripted IGOR procedures. The first AP evoked by the square pulse protocol was used to calculate the AP properties. The AP threshold was defined as the minimum of the first derivative of the AP (= the point of inflection during the depolarization). The afterhyperpolarization is the minimum after the AP peak. The amplitude is measured between RMP and AP peak. To calculate the AP half-width, the half distance between threshold and peak is measured and the distance between this point during depolarization and repolarization is evaluated. The time to peak is the duration between current pulse onset and AP peak. The maximum slope of the upstroke was calculated between threshold and peak, whereas the slope of the subthreshold depolarization was determined between RMP and threshold. Overshoot slope was calculated for the part exceeding 0 mV.

Sine wave affinity (sine_score) was calculated as the difference in the number of spikes between 20 sinusoidal and 20 rectangular stimulations (theoretical maximum = 20 (20 sine spikes vs 0 rectangular spikes), theoretical minimum = -20 (0 sine spikes vs 20 rectangular spikes).

Following frequency (FF) is given as the number of spikes elicited by 20 stimulations for each stimulation frequency.

Half Sine Threshold was defined as amount of current injection in which the AP was elicited during half sine stimulation.

Activity dependent Time to Peak delay was extracted as the change in time to peak in milliseconds compared to the first AP evoked by 75 suprathreshold stimulations. Activity dependent change in resting membrane potential is given as change in mV of the membrane potential during stimulation compared to potential before stimulation by 75 suprathreshold square voltage pulses. Activity dependent change in slope is given as maximum in first deviation of AP train normalized to value of first AP evoked by 75 stimulations.

#### RNA-Sequencing

##### snRNAseq dataset

###### Construction of single-nucleus gene expression libraries and sequencing

At the next generation sequencing core facility of the Medical Faculty Mannheim at Heidelberg University, single-nucleus solutions were subjected to barcoding, reverse transcription and gene expression library construction using the 10XGenomics microfluidics platform (Chromium controller and Next GEM Single Cell 3’ v3.1 (sample 1-3) or of Multiome Kit (sample 4-5)) according to the manufacturer’s instructions. Gene expression libraries were sequenced on an Illumina NextSeq550.

###### snRNAseq data analysis

Sequencer output files were converted into fastq-files using Cellranger 6.0.1 mkfastq. Using Cellranger 6.0.1 count, reads were then mapped to the pig genome (Sscrofa11.1 with Ensembl 105 annotation) on the bwForcluster Helix high-performance computing system. Reads uniquely mapping in sense to the transcriptome (exon and intron) were then used to quantify transcript abundances using unique molecular identifiers (UMI). After cell calling, which is also built into Cellranger 6.0.1, a cell-gene expression matrix was generated and imported into R (version 4.2.2, RStudio version 2022.12.0) for further analysis using Seurat (version 4.3.0.9001). Nuclei with less than 200 detected UMIs and doublets (identified with DoubletFinder) were filtered out. For each replicate, mitochondrial reads were regressed out during count normalization with “SCTransform” v2. Normalized data was then used for anchor-based data integration resulting in an integrated data assay. This data assay (variable features set to 3,000) was used for dimensional reduction by principal component analysis. The first 30 principal components were used to visualize nuclei in two dimensions using uniform manifold approximation and projection (UMAP). Similarly, the first 30 principal components were used for cluster analysis setting the resolution parameter of “FindClusters” to 2. Clusters with high percentage of mitochondrial genes or low number of DEGs likely reflect empty droplets and were therefore removed from the dataset. The resulting dataset (16,979 neuronal and non-neuronal nuclei) was subset to neurons based on the expression of SNAP25, SCN9A, THY1, TAC1, and RBFOX1 (pan-neuronal markers of human DRG neurons (*27*)). A neuronal set of 3,000 variable features was identified and dimensional reduction, two-dimensional projection and cluster analysis was repeated with the same parameters. This resulted in 18 clusters, three of which were merged based on similar marker gene expression. Finally, annotation of the 16 final clusters was performed based on the presence/absence of marker genes used to distinguish transcriptionally-defined sensory neurons in mice (*30, 39, 63–66*), monkey (*25, 26*) and human (*25, 27, 29*).

##### Patch-seq Dataset

###### Sequencing and pre-processing

Individual lysed cells were treated according to the Smart-Seq2 library preparation protocol (*67*). Preamplified cDNA was quantified and the average size distribution was determined via D5000 assay on a TapeStation 4200 system (Agilent). Tagmentation and subsequent next-generation sequencing (NGS) library generation were performed using 200 pg of cDNA. NGS libraries were quantified by High-Sensitivity dsDNA assay on a Qubit (Invitrogen) and the average size distribution was determined via D5000 assay on a TapeStation 4200 system (Agilent). Libraries were equimolarly pooled, and sequenced SR 75bp on a NextSeq500 system with a 75 cycles High Output v2 chemistry or NextSeq2000 with a P2 or P3 100 cycles flow cell (Illumina). Raw sequencing data were demultiplexed and converted into fastq format using bcl2fastq2 v2.20.

After quality checking with MultiQC v.1.5, reads were pseudoaligned to the Sus Scrofa genome 11.1 (104, Ensembl) via Kalisto v.0.440 with default parameters. Patch-seq cell samples with fewer than 10,000 reads were discarded.

#### Spatial Transcriptomics Dataset

##### Fixation, Staining and Imaging

Methanol fixation and Eosin and Haematoxylin staining for an initial tissue quality control analysis was performed as described in the 10XGenomics Methanol Fixation, H&E Staining & Imaging for Visium Spatial Protocols (Demonstrated Protocol CG000160). We stained a total of 60 sections, each 10 µm in thickness, from 22 fresh frozen DRG obtained from 7 female pigs.

These sections were mounted within the capture areas of 11 Visium slides that contained 55-μm printed barcoded spots. Imaging was conducted using the manual load and the fluorescence features of an Olympus vs120 Slide Scanner.

##### Tissue Optimization

Different permeabilization times (3, 6, 12, 18, 24 and 30 minutes) were evaluated following the 10XGenomics Visium Spatial Gene Expression Reagent Kits - Tissue Optimization User Guide CG000238 Rev E. This protocol required the reagents from the Visium Spatial Tissue Optimization Reagent Kit, PN-1000192 (stored at −20°C) and the Visium Spatial Tissue Optimization Slide Kit PN-1000191 (stored at ambient temperature). The exposure time of the permeabilization enzyme was 18 minutes based on the analysis of the tissue optimization experiment

##### 10XGenomics Visium Spatial protocols

We used the spatial sequencing protocol Visium Spatial Gene Expression Reagent Kits, 16 rxns PN-1000186 and Library Construction Kits, 16 rxns PN-1000190. All reagents were stored at - 20°C. Additionally, Visium Spatial Gene Expression Slide Kits, 16 rxns PN-1000185 were acquired and stored at ambient temperature.

The protocol was performed exactly as stated in the Visium Spatial Gene Expression Reagents Kits User Guide CG000239 Rev F. It consisted of a 5-step process, starting with permeabilization and reverse transcription (step 1). The exposure time of the permeabilization enzyme was 12 minutes based on the analysis of the tissue optimization experiment. After second strand synthesis and denaturation (Step 2), the samples were prepared for a full-length cDNA amplification via Polymerase Chain Reaction (PCR) (step 3). Afterwards, a Visium spatial gene expression library was generated (step 4), followed by sequencing (step 5). Steps 4 and 5 were performed at the Genomics Core facilities of the University of Texas at Dallas. The samples were sequenced using the Illumina NextSeq2000 Sequencing system.

##### Visium spatial RNA-seq analysis

From our sequencing run, we obtained a total average of 59,081,551 M reads, 86.51% of the reads were mapped with 84.22% confidence. We detected an average of 13,870 genes and 1574 barcodes under tissue. The mean of reads per barcode was 40,490, and the median of Unique Molecule Identifier (UMI) counts per spot was 2,819. The generated Illumina BCL files were processed with the 10XGenomics pipeline (Space Ranger v1.1). This pipeline allowed the alignment of the FASTQ files with bright-field microscope images and the pig reference transcriptome (Sscofra11.1). Using the Loupe Browser (v4.2.0, 10x Genomics), we visualized the location of the barcoded mRNAs for downstream analysis by selecting the barcoded spots overlapping single neurons. We identified 6408 barcodes overlapping single neurons and 12,304 overlapping multiple barcodes. To avoid double counts, we excluded the barcodes overlapping multiple neurons. As part of a quality control step, we removed neuronal barcodes with less than a hundred reads and no counts of the neuronal marker Synaptosome Associated Protein 25 (*SNAP25*) using Python (v3.8 with Anaconda distribution).

##### Visium spatial morphology measurements

We compiled a .csv file containing the positional data (coordinates) of barcodes overlapping with single neurons per Visium capture frame. Subsequently, we employed Cellsens imaging software to visualize the corresponding .tif image files and identify the neurons. Within this software, we utilized the polyline measuring tool to determine the diameter of neurons (µm) exhibiting distinct DAPI nuclear staining, indicating their sectioning within the central region of the cell. Cells displaying cytoplasmic distortion attributed to freezing artifacts were excluded from our measurements.

#### RNA-seq dataset integration

##### Pig cross-dataset genomics dataset integration and cell type mapping

We used the pig snRNAseq dataset as a reference to integrate each of our additional pig datasets separately, ensuring precise cell-type mapping across species and technologies. The datasets integrated with this reference include Patch-seq data from pigs, and a separate 10X Visium spatial transcriptomics dataset from pigs. In our integration process, the Patch-seq dataset featured 22,259 genes and the pig 10X Visium had 15,924 genes. To align these with our pig snRNAseq reference, which contains 30,477 genes, we used the intersection of the gene symbols across each pair of datasets. To integrate Patch-seq transcriptomes with the snRNAseq reference dataset, we used the Canonical Correlation Analysis (CCA) workflow from Seurat (v4.3.0). We performed dataset integration by treating the Patch-seq data as coming from a single separate batch, re-integrated all samples across all batches, using SCTransform v2 normalization with mitochondrial reads regressed out. We selected 6,000 highly variable genes as anchor integration features to integrate data across batches into a single integrated atlas object using the SelectIntegrationFeatures function. The datasets were then integrated using the IntegrateData function with the ‘normalization.method’ parameter set to “SCT”, and ‘k.weight’ set to 75. Next, we performed principal component analysis (50 dimensions), and UMAP on the first 30 principal components for dimensionality reduction and visualization purposes.

To assign cell-type labels to the Patch-seq sampled cellular transcriptomes, we used a K-nearest neighbour approach based on the 20 nearest reference snRNAseq neighbours using the first 30 PCA dimensions. The nearest neighbours for each Patch-seq cell were first defined using FindNeighbors in Seurat, then identified using the function TopNeighbors with *k* parameter of 20. Each Patch-seq neuron’s cell type was defined as the most frequent snRNAseq-based cell type present in the nearest 20 neighbours.

We used a modified approach of the above for integrating the pig 10X Visium dataset and to map assigned cell-type labels, due to the larger size of the 10X Visium datasets relative to the snRNAseq data. For the 10X Visium dataset, we iterated through batches of a maximum size of 250 cells within each donor to facilitate using the same parameters for the integration and cell type mapping procedure as were used for integration and cell-type mapping of the Patch-seq data.

##### Analysis of species and dataset-specific transcription profiles

To identify transcriptional markers of each DRG neuron subtype, we used the FindAllMarkers() function from Seurat (v4.3.0) separately on each collected dataset. We set the parameters min.pct and logfc.threshold to -Inf, min.cells.feature and min.cells.group to 1 to maximize the number of genes tested for comparison across the datasets.

##### Transcriptome marker gene correlation analyses

To determine agreement of transcriptional markers across the datasets, for each dataset, we used the output of the FindAllMarkers() function from Seurat to compute enrichment fold changes (avg_log2FC) of each gene from each cell-type in comparison to all other cells in the dataset.

Next, we intersected the lists of enriched genes from each cell type between pairs of datasets and correlated enrichment values across intersecting genes. We then used the cor.test() function in R with the method parameter set to "spearman" to correlate the avg_log2FC values for each gene across the two datasets.

##### Principal Component Analysis (PCA) of Electrophysiological Features

Electrophysiological data from patch-clamp recordings were extracted for four key features associated with CMi-like properties: action potential duration (APD), response to 50 Hz stimulation (50Hz), sinus score, and relative activity-dependent slowing (Rel. ADS). To address missing values in the dataset, we employed k-nearest neighbor (kNN) imputation using the recipes package in R. The imputation step used 5 neighbors (k = 5) to estimate missing values based on similar cells in the multidimensional feature space. This approach preserves the overall structure of the data while allowing for complete case analysis in subsequent steps.

Following imputation, the four electrophysiological features were scaled and centered. We then performed Principal Component Analysis (PCA) using the prcomp function in R. The first two principal components were used for further analysis and visualization. We created a biplot to display both the distribution of cells in the PC space and the contributions of each electrophysiological feature to the principal components. Cell types were color-coded, and feature loadings were displayed as vectors on this plot.

##### CMi associated differential expression analysis

To identify genes associated with CMi-like properties, we performed differential expression analysis using the limma-voom pipeline. The CMi score (PC1 from the electrophysiological PCA) was used as a continuous variable in the design matrix. Genes with an adjusted p-value < 0.05 and ABS(log2 fold change) > 0.5 were considered significantly differentially expressed.

##### Cross-species cell type replicability and prediction

We compared our pig snRNA-seq dataset to the cross-species DRG atlas from Bhuiyan et al. (2024) using two complementary approaches: MetaNeighbor for cell type replicability assessment and Seurat’s label transfer for cell type prediction.

##### Metaneighbor analysis

MetaNeighbor (version 1.16.0) was used to assess cell-type replicability between our pig snRNA-seq data and the cross-species atlas. MetaNeighbor constructs a network of rank correlations among all cells based on shared variable genes, using the principle that cells belonging to the same subtype exhibit more correlated gene expression patterns than distinct cell types. By applying a neighbour voting system, each cell accrues a score denoting the proportion of its neighbours sharing its cell type. This metric is quantified by the area under the receiver operator characteristic curve (AUROC), with the mean AUROC across folds determining the degree of cell type similarity. The unsupervised procedure requires a gene-by-cell matrix in SummarizedExperiment format, along with metadata identifying dataset and cell-type labels for each cell. We followed the unsupervised MetaNeighbor procedure described in detail with accompanying scripts labelled Procedure 1 at https://github.com/gillislab/MetaNeighbor-Protocol/ (*68*). We identified 1172 highly variable genes using the variableGenes() function.

These genes were input for MetaNeighborUS() to generate a cell-cell similarity network and quantify cell type replicability. The output is a cell type by cell type mean AUROC matrix, with higher values indicating greater similarity. We used MetaNeighborUS() with parameters one_vs_best = TRUE and symmetric_output = FALSE to identify the best matches between pig and atlas cell types. A cluster graph was created using makeClusterGraph() (low_threshold = 0.3) and visualized with plotClusterGraph() to represent cell type relationships across the datasets.

##### Label transfer analysis

To predict cell types for our pig snRNA-seq data, we used Seurat’s label transfer workflow (version 4.3.0.1), employing the same cross-species DRG atlas as a reference using its ‘Atlas_annotation’ cell-type labels. The pig snRNA-seq data was preprocessed using standard Seurat functions (NormalizeData, FindVariableFeatures, ScaleData, RunPCA). We used FindTransferAnchors() to identify anchors between the atlas and the pig data, specifying "RNA" as the assay and "cca" as the reduction method. TransferData() was then applied to transfer cell type labels from the atlas to the pig dataset. Finally, we used MapQuery() to project the pig data onto the UMAP space of the atlas for visualization and comparison of predicted cell types.

### Computational action potential modeling

We used computational modeling to study the impact of SCN11A abundance on the AP morphology. We adapted the modeling framework from (*43*) to C-OSMR-SST neurons. The model considers SCN1A-5A & 8A-11A and is based on an extension of the Hodgkin-Huxley approach which accounts for non-exponential inactivation kinetics and shows better agreement to data. The gating parameters were fitted based on voltage clamping data and are taken from (*43*). The contribution of a given sodium channel subtype to the total sodium conductance is chosen according to its expression level in C-OSMR-SST neurons. The Aps were generated by a current injection of 30µA/cm^2 starting at time 0 and lasting until the end of the observation period. The AP width was measured at the threshold voltage which was determined using the second derivative of dV/dt with respect to V as described in (*69*).

### In vivo experiments

#### Pig

All experimental interventions in pigs were approved by the regional ethics council in Karlsruhe, Baden-Wuerttemberg, Germany (G-78/18). *In vivo* extracellular recordings from the “teased” saphenous nerves were performed using DAPSYS software (www.dapsys.net) (*70*). Briefly, electrical rectangular pulses (0.5 ms duration; 20 mA intensity; 0.25 Hz) were delivered by a constant current stimulator (DS7A, Digitimer Ltd., Hertfordshire, UK) via two non-insulated microneurography electrodes (FHC Inc., Bowdoin, ME, USA), inserted intradermally at sites where time-locked action potentials with long latencies could be elicited (*22*). At the end of the experiment, pigs were euthanized by i.v. injection of 10 ml Tanax (T-61, Intervet Deutschland GmbH) and death was confirmed by induction of lasting electrical silence on ECG and disappearance of carotid pulse.

#### Human

##### OSM experiments

Recombinant Human Oncostatin-M protein, carrier free (Biotechne, Minneapolis, Minnesota, U.S., REF Number: 8475-OM-050/CF), diluted in 20 µl sterile synthetic interstitial fluid (SIF) (*71*) was injected intracutaneously into human volunteers, who provided informed consent. The experiments were conducted on independent researchers who are all senior co-authors of this study (self-experiments). SIF contained (in mM) 107.8 NaCl, 3.5 KCl, 1.5 CaCl2, 0.7 MgSO4, 26.2 NaHCO3, 1.7 NaH2PO4, 9.6 sodium gluconate, 5.5 glucose, and 7.7 sucrose with a stable pH of 7.4. The test person was blinded towards the applied substances during experiments.

##### Superficial blood flow

Superficial blood flow was assessed by a laser Doppler imager (LDI, Moor Instruments Ltd., Devon, United Kingdom) or a laser speckle imager (FLIPI2, Moor Instruments Ltd., Devon, United Kingdom with measurement software V2.0). Baseline images were taken before and 24 h after the injections. Erythema areas were calculated as those pixels exceeding the mean flux value + 2-fold standard deviation measured in a control area between the injection sites using moorLDI Software (Research Version 5.3, 2009, Moor Instruments Ltd) and for laser speckle data moorFLPI-2 software (Review Software V 5.0, Moor Instruments Ltd)).

##### Experimental protocol for psychophysics

10 µl in SIF diluted OSM and SIF alone as control were intracutaneously injected at the volar forearms of healthy volunteers in a blinded approach. Thereafter ratings were assessed verbally every 10 seconds for 5 minutes on a rating scale from 0 (no pain/itch) to 10 (maximally imaginable itch/pain) and skin vasodilation was measured using either laser Doppler or laser speckle imaging as described above. Thereafter, and 24 h after skin injection, vasodilation via Laser speckle and laser Doppler imaging and hyperalgesia were assessed using manual pressure, mechanical impact and von Frey filament stimulation.

##### Microneurography

Experiments were conducted at the University of Aachen, microneurography protocols were approved by the local ethics committee (EK 143/21) and are registered within the German Registry for Clinical Studies (DRKS00025261). APs of single C-fibers from cutaneous C-fiber fascicles of the superficial peroneal nerve were recorded as previously described (*10, 72*).

A tungsten recording needle (Frederick-Haer, Bowdoinham, ME, USA) is inserted and placed close to an unmyelinated afferent nerve fiber bundle. After reaching a stable position, C-unit innervation territories are detected using a pointed electrode (0.5 mm diameter) delivering electrical pulses.

C-fiber units are identified by their low conduction velocity (< 2 m/s). A pair of 0.2 mm diameter needle electrodes (Frederick-Haer company) is inserted into the previously located innervation territory for intracutaneous stimulation of the recorded C-fibres. Low repetition rates are inserted using a Digitimer DS7 constant current stimulator. The signal is amplified, filtered and stored on a computer using custom-written microneurography software DAPSYS and analyzed offline using DAPSYS (Brian Turnquist, http://dapsys.net) and Microsoft Excel.

Single C-fibers were differentiated by their individual conduction latency during continuous low frequency stimulation (0.25 Hz; intensity at least 1.5 times the individual electrical fiber’s threshold). After recording C-fiber responses, we used the “marking technique” to characterize the units. This is based on the slowing of conduction velocity when a C-fiber conducts more than one AP within a short time period, which is known as activity-dependent conduction velocity slowing (ADS). The amount of ADS strongly correlates with the number of additional APs conducted in the seconds before the electrically induced AP.

To determine the mechanical sensitivity of the recorded C-fibers, we repetitively applied mechanical stimuli using stiff von Frey filaments of 22 g (Stoelting, Chicago, IL, USA) in the receptive fields.

We assigned C-fibers as mechanosensitive (CM) or mechano-insensitive (CMi) according to their mechanical responses and electrophysiological properties. C-fibers with an ADS < 5% of their initial latency to an electrical stimulation protocol with rising frequencies (20 pulses at 0.125 Hz, 20 pulses at 0.25 Hz, 30 pulses at 0.5 Hz), a normalization of latency thereafter of more than 43% within 20 s, and a response to < 22 g von Frey stimulation, were classified as CM-fibers. C-fibers with an ADS > 5% and a recovery of < 43% and no mechanical response to 75 g von Frey stimulation were classified as CMi-fibers.

##### Experimental protocol for microneurography

First the C-fibers were classified into CM and CMi as described above. In 50 healthy participants 2 Hz stimulation was performed for 3 minutes. Part of those data were utilized for comparison to patch clamp data (see figure 3). In the self-experiments in a different experimental setup during continuous 0.25 Hz electrical stimulation of the fibers in the receptive field in the skin 20 µl of OSM was injected intracutaneously directly under the stimulation needles. Thus, OSM affected maximally 5 mm of the nerve fibers. Continuous electrical stimulation is necessary for using the marking method for assessing fiber activation and changes in biophysical properties (*72*).

### Statistics

All statistical analyses were conducted using R (version 4.2.1). Non-parametric tests, including the Wilcoxon Rank-Sum Test and the Mann-Whitney U Test, were applied to evaluate median differences in data sets that were not assumed to follow a normal distribution. The Kruskal-Wallis Test was used for comparing medians across multiple groups, while the Welch’s t-test was used to compare means between two groups with unequal variances. For analyses involving multiple group comparisons, ANOVA was used to ascertain mean differences, followed by Tukey’s Honest Significant Difference (HSD) Test for post-hoc pairwise comparisons to control for multiple testing errors. Specifics regarding the number of observations, p-values, and the tests used are reported in the figure legends or directly in the main text as appropriate.

Principal component analysis was performed with Graphpad PRISM (version 10.2.0) on standardized data. Principal components were included that together explained >75% of the total variance.

## Acknowledgments

The authors thank the FlowCore and next generation sequencing core facilities of the Medical Faculty Mannheim at Heidelberg University for their assistance with nuclei sorting and generation of single-nucleus gene expression libraries and sequencing. We also thank Heidi Theis (PRECISE, DZNE, Bonn) for preparation of Patch-seq libraries. The authors thank the organ donors and their families for their enduring gift. The authors thank the Genome Center at The University of Texas at Dallas for the services to support pig DRG spatial sequencing.

## Funding

Clinician Scientist program of the Faculty of Medicine of the RWTH Aachen university (JK)

DFG, German Research Foundation 363055819/GRK2415 (AL)

DFG, German Research Foundation 368482240/GRK2416 (AL)

DFG, German Research Foundation LA 2740/6-1 (AL)

DFG, German Research Foundation NA 970/5-1 (BN)

DFG, German Research Foundation 255156212/CRC1158/Z02-INF (HJS)

DFG, German Research Foundation 350193106/FOR2690/TP01 (HJS)

DFG, German Research Foundation 255156212/SFB1158 A01 (MS)

DFG, German Research Foundation 350193106/FOR2690 (MS)

DFG, German Research Foundation 1587/10-1 (IK)

DFG, German Research Foundation 1587/11-1 (IK)

Interdisciplinary Center for Clinical Research within the Faculty of Medicine at the RWTH Aachen University IZKF TN1-1/IA 532001 (AL)

CAMH Discovery Fund, Krembil Foundation, Kavli Foundation, McLaughlin Foundation, Natural Sciences and Engineering Research Council of Canada (RGPIN-2020-05834 and DGECR-2020-00048 (DH,ST)

Canadian Institutes of Health Research (NGN-171423 and PJT-175254) (DH,ST)

Simons Foundation Autism Research Initiative (DH,ST)

Baden-Württemberg through bwHPC and the German Research Foundation (DFG) through grant INST 35/1597-1 FUGG (HJS, MS)

NIH grant U19NS130608 (TJP)

German Research Foundation (DFG) through grant INST 35/1503-1 FUGG (HJS,MS)

Interdisciplinary Center for Clinical

Research within the faculty of Medicine at the RWTH Aachen University (BN)

NIH U19 NS130617, Rita Allen Foundation, Burroughs Wellcome Fund Career Award in Medical Sciences (WR)

## Author contributions

Conceptualization: JK, DH, HJS, TJP, MS, BN, ST, AL.

Methodology: JK, DH, HJS, MMM, NH, IT, MB, BN, ST, AL, TS.

Software: JK, DH, MMM, DT, NNI, TS.

Validation: JK, DH, HJS, ST, AL.

Formal analysis: JK, DH, HJS, MMM, AF, DT, AM, SAB, BN.

Investigation: JK, HJS, MMM, NH, AF, IT, RB, IS, DT, SS, AM, LB, JSS, IK, MS, BN, ST, TS.

Resources: DT, LE, MB, BN, AL.

Data Curation: JK, DH, HJS, MMM, AF, AM, MB, BN, ST, AL.

Writing - Original Draft: JK, DH, MMM, TJP, ST, AL.

Writing - Review & Editing: JK, DH, HJS, MMM, NH, AF, AM, LE, MB, IK, MS, SAB, WR, BN, ST, AL.

Visualization: JK, DH, HJS, MMM, NH, ST.

Supervision: MB, TJP, BN, ST, AL.

Project administration: JK, HJS, MB, NH, TJP, ST, AL.

Funding acquisition: JK, HJS, IK, WR, TJP, BN, AL.

## Data and materials availability

All raw gene expression data with detailed metadata supporting the findings of this study will be openly after peer-reviewed publication.

Integration and analysis code, as well as extracted patch recording features will be available after peer-reviewed publication.

---

**Figure S1:**
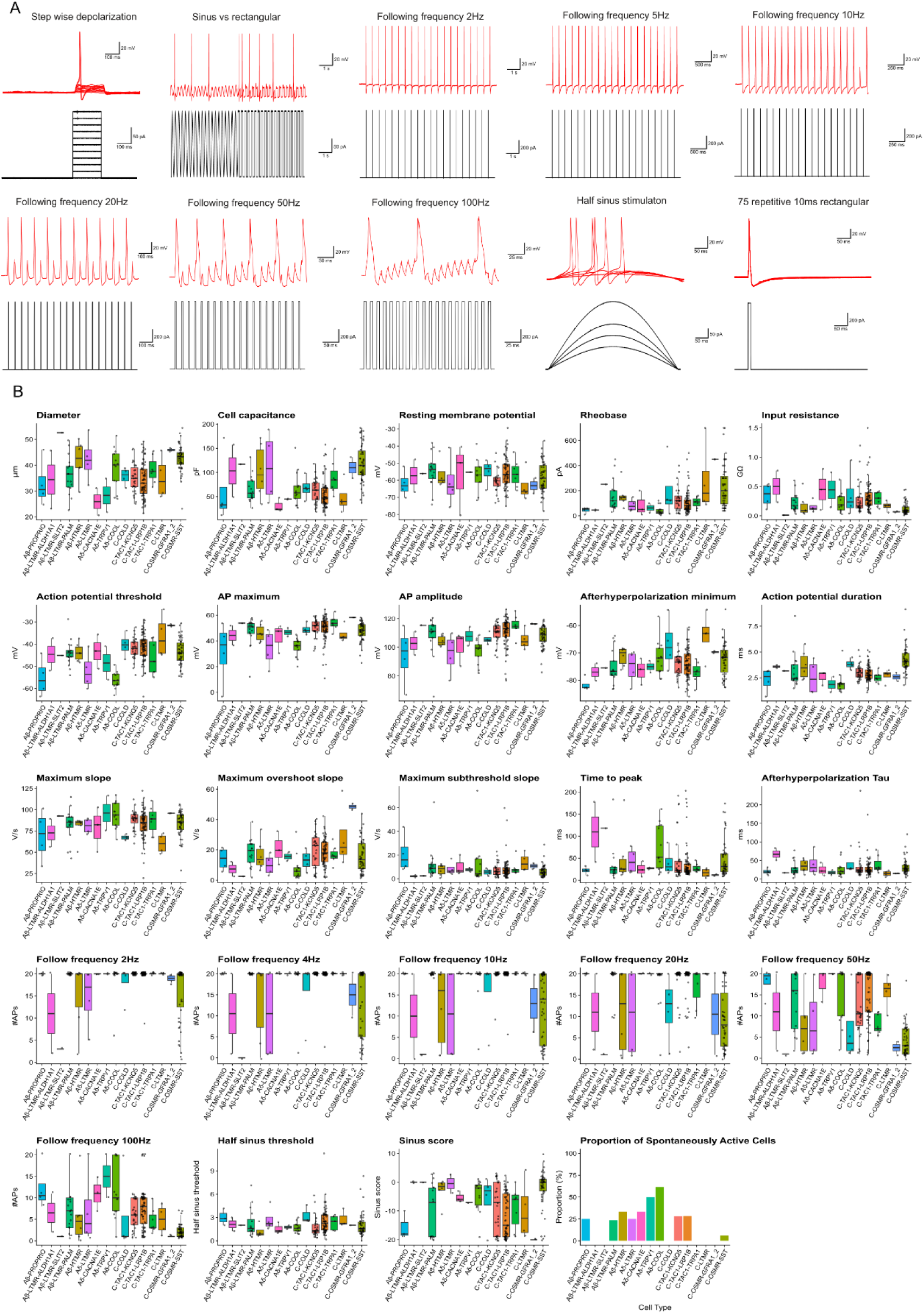
Extended electrophysiological data. **A**) Pulse protocols (black) for electrophysiological assessment and example patch-clamp recording traces of pig DRGs (red). From up left to down right: 200 ms stepwise depolarization protocol, sine vs rectangular 4 Hz stimulation, 20 1 nA pulses at 2, 5, 10, 20, 50 and 100 Hz, 500 ms half sine-shaped stimulation, 75 repetitive 1 nA 10 ms stimulations. **B**) Combined scatter and boxplots showing quantification of each electrophysiological feature per mapped transcriptomic identity.

**Figure S2:**
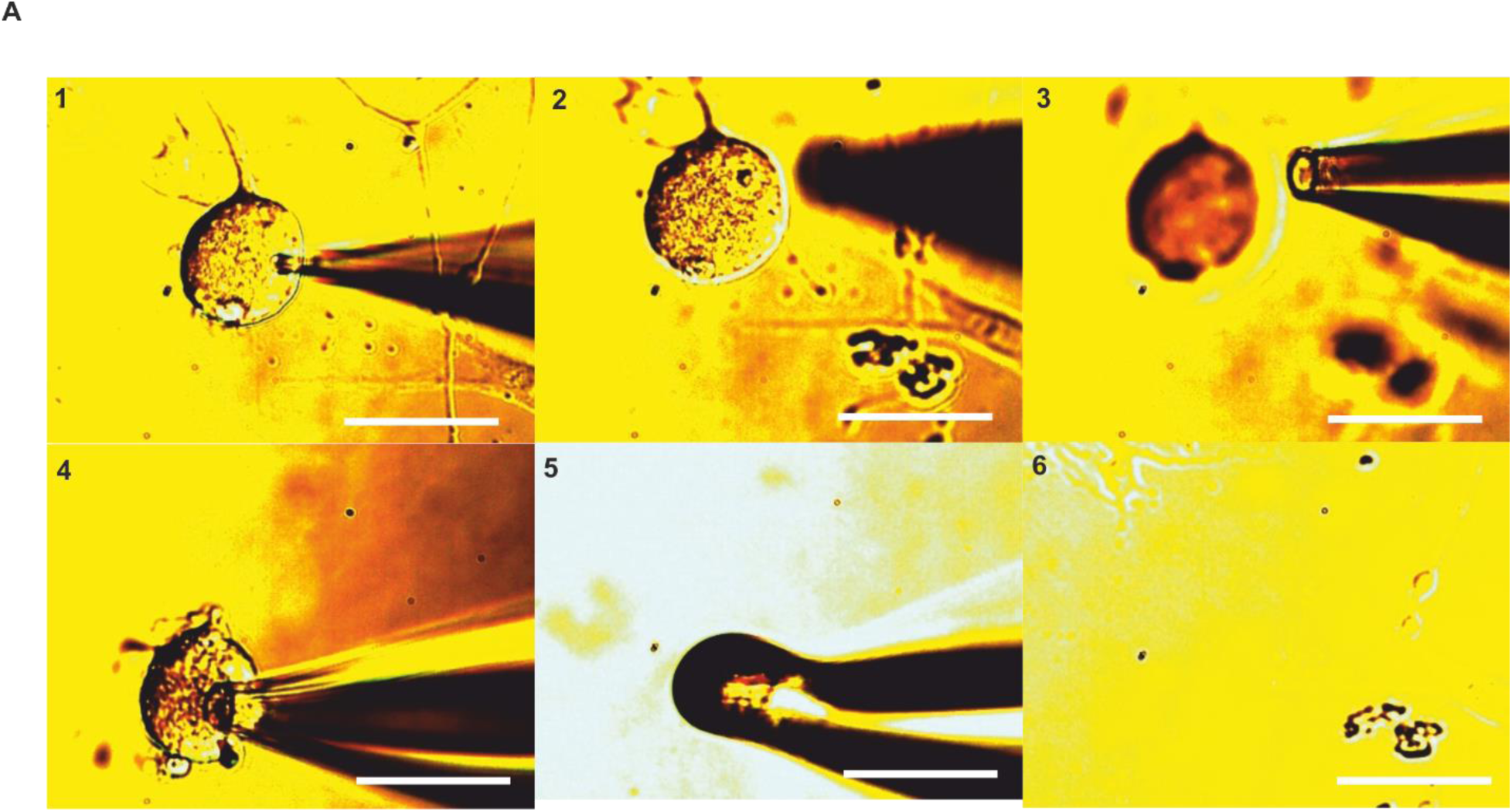
Patch-seq whole neuron sampling procedure. Light microscopy time-sequence of one cultured DRG neuron being harvested for Patch-seq following electrophysiological characterization. Scale bars indicate 50 µm. Up left to down right: 1) Neuron shown with small diameter pipette for patch clamp based current clamp recordings. 2) After recording, the recording pipette is detached, and cytosolic content washed in the pipette tip is transferred to a PCR tube containing lysis buffer. 3) A second, larger diameter pipette is used for extraction of the entire neuron with neurites. 4) The second pipette is attached to the neuron using strong negative pressure and the neuron with its neurites is carefully detached and extracted from the coverslip. 5) Image of the second pipette with the neuron attached shown outside (i.e., above) the extracellular solution, following which the neuron is carefully placed into the same PCR tube from step 2 containing lysis buffer. 6) Image of the former position of the neuron, confirming completion of neuronal harvest.

**Figure S3:**
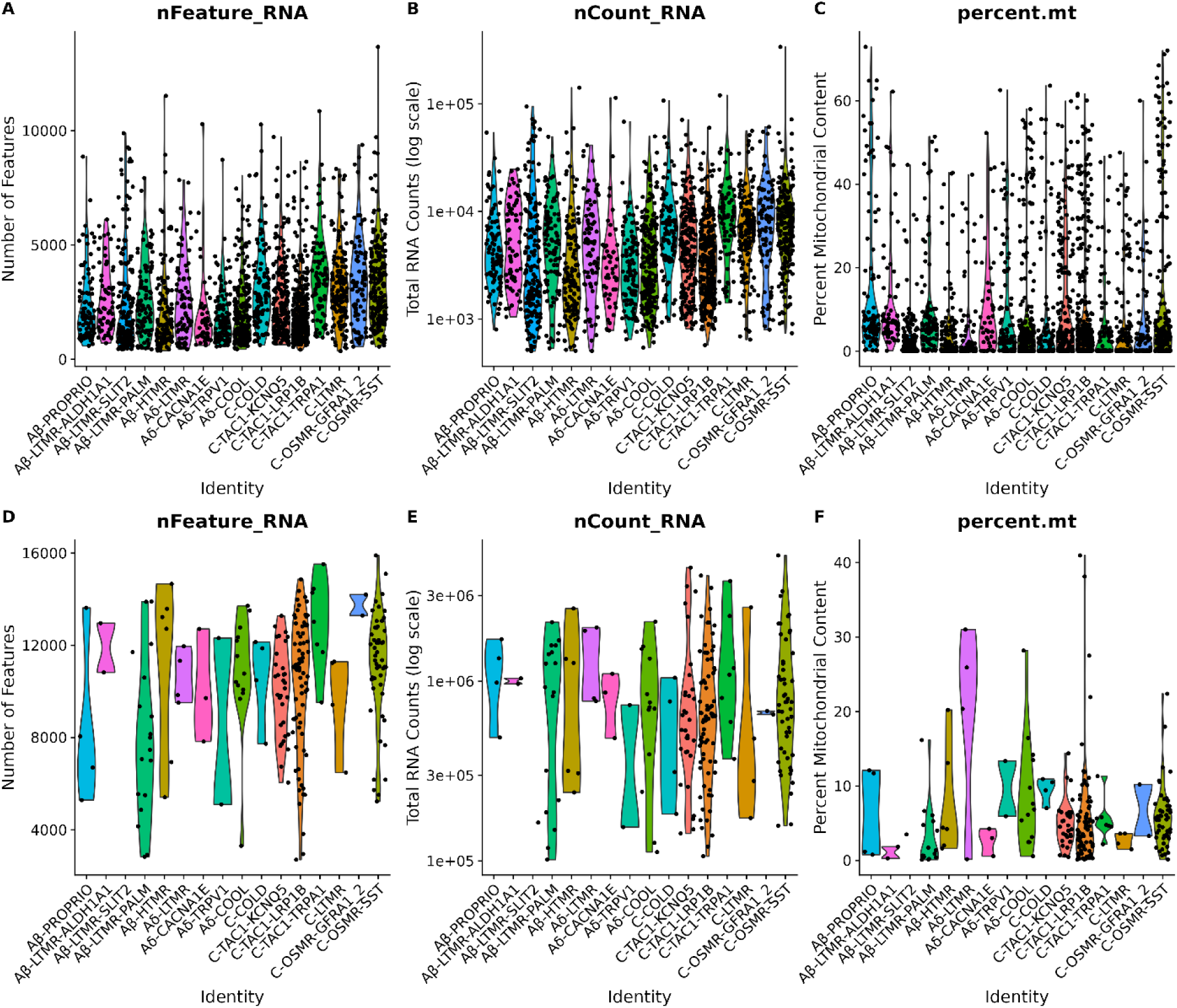
Sequencing metrics of snRNAseq and Patch-seq datasets. **A-C**) Sequencing metrics for the snRNAseq dataset, per assigned neuronal identity: a) number of genes per cell, b) number of reads per cell, c) fraction of mitochondrial reads. **D-F**) Corresponding sequencing metrics for the Patch-seq dataset: d) number of genes per cell, e) number of reads per cell, f) fraction of mitochondrial reads.

**Figure S4:**
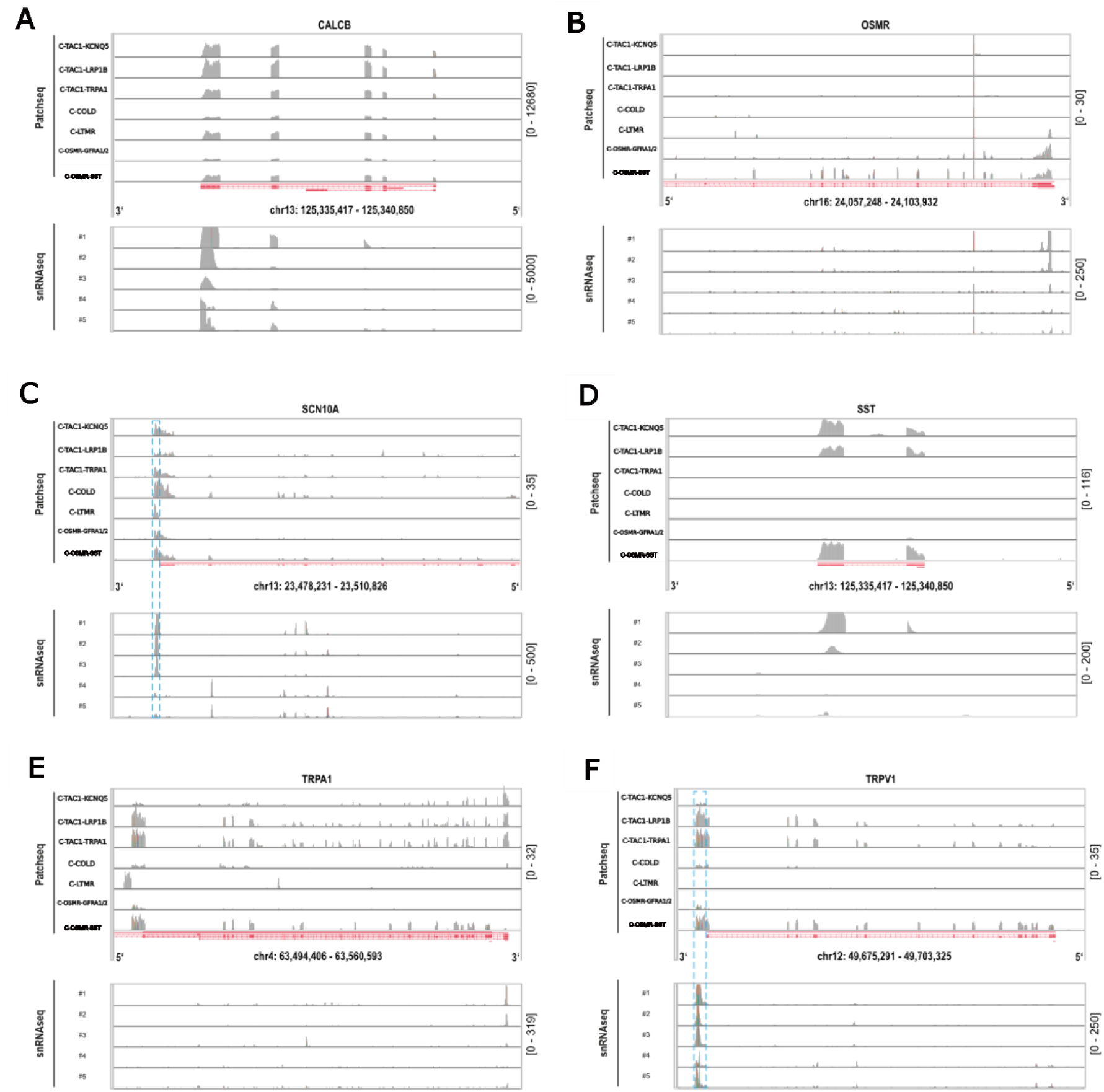
Illustration of how limitations in porcine reference genome annotation likely impact gene expression quantification. **A-F**) Read coverage plots for exemplar genes from Patch-seq (one representative cell per C-fiber type, top) and snRNAseq (merged data from multiple cells separated by barcoding reaction, bottom) datasets. X-axis denotes genome position (3’ and 5’ gene ends annotated per gene), and y-axis denotes read coverage (scales shown on right). Red track in middle illustrates gene models from pig reference transcriptome annotation (Ensembl 105). Note that Smart-seq2 protocol, used for reverse transcriptase for Patch-seq, results in full-length cDNA generation and thus more complete coverage of mRNA throughout the whole gene body. In contrast, given our usage of the 10XGenomics Single Cell 3’ Kit, as expected, snRNAseq read coverage are biased towards 3’ ends of gene bodies. We note that this qualitative inspection revealed that some genes, including SCNA10A (c) and TRPV1 (f), show some evidence of having several unannotated reads in the 3’ UTRs of these genes (highlighted by blue dashed lines), which we reason is likely due to incomplete annotation of the 3’ UTRs of these genes in the used reference transcriptome. However, this issue appears to be gene specific as the other genes manually assessed, including CALCB (a), OSMR (b), SST (d) do not appear to qualitatively display evidence of this issue. Note that while 3’ UTR appears well annotated for TRPA1 (e), 5’ UTR appears potentially misannotated, as our Patch-seq datasets showed evidence that some reads were unannotated in these regions.

**Figure S5:**
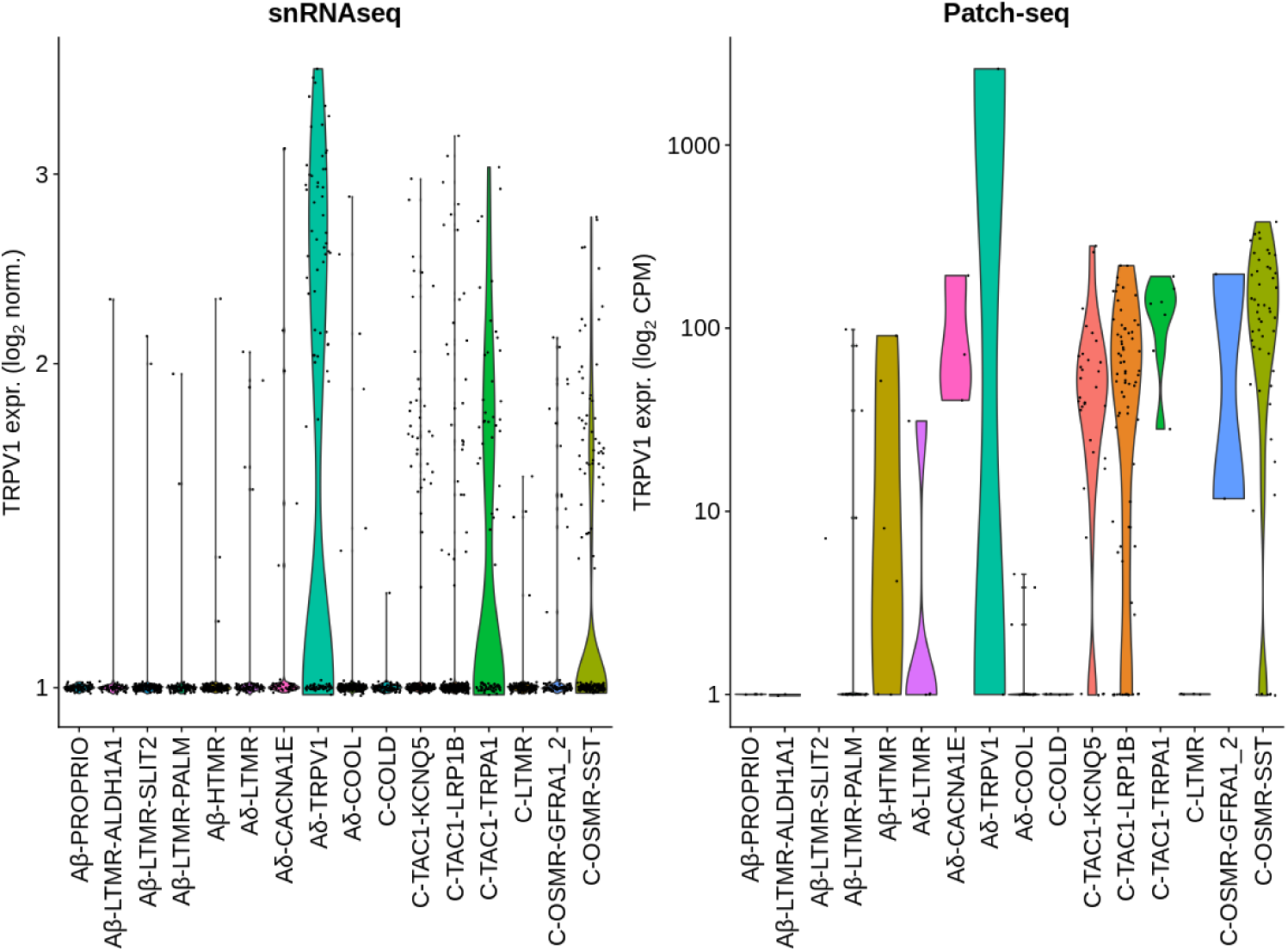
TRPV1 expression is consistent between snRNAseq and Patch-seq datasets. Violin plots showing TRPV1 expression levels across neuronal subtypes in snRNAseq (left) and Patch-seq (right) datasets. The three clusters with highest TRPV1 expression (Aδ-TRPV1, C-TAC1-TRPA1, and C-OSMR-SST) are consistently identified across both technologies. Y-axis shows normalized expression values (log2 normalized counts for snRNAseq; log2 CPM for Patch-seq). Each dot represents an individual cell/nucleus.

**Figure S6:**
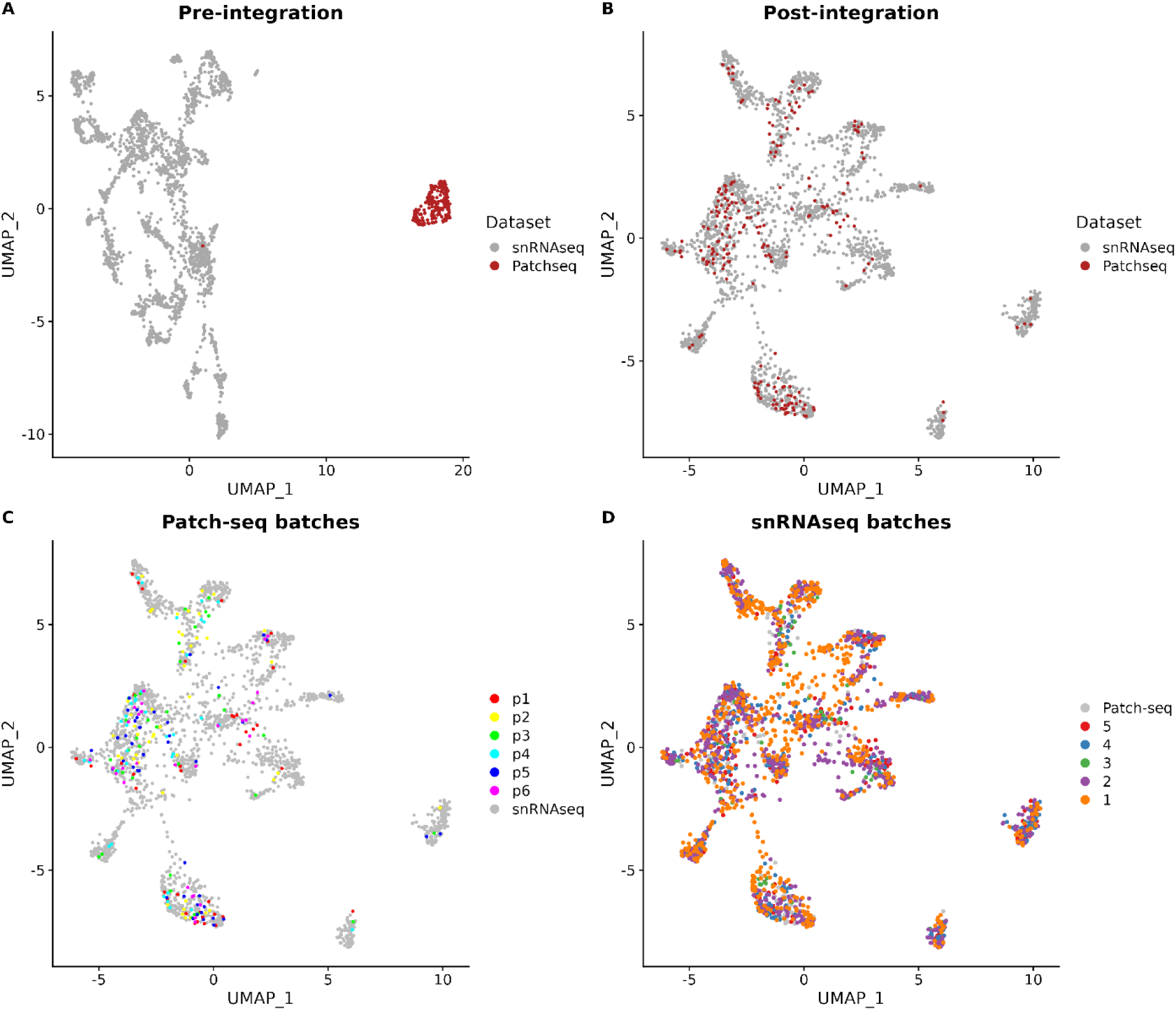
Integration of snRNAseq and Patch-seq datasets and batch distribution across UMAP space. UMAP visualizations of single-cell transcriptomics data from pig dorsal root ganglia (DRG) neurons. (A) Pre-integration view showing separate clusters for snRNAseq (grey) and Patch-seq (red) datasets. (B) Post-integration view demonstrating successful merging of snRNAseq and Patch-seq data. (C) Integrated data colored by individual Patch-seq experimental batches (p1-p6), with snRNAseq cells shown in grey. (D) Integrated data colored by individual snRNAseq animal donors (1-5), with Patch-seq cells shown in grey. Each point represents a single cell or nucleus.

**Figure S7:**
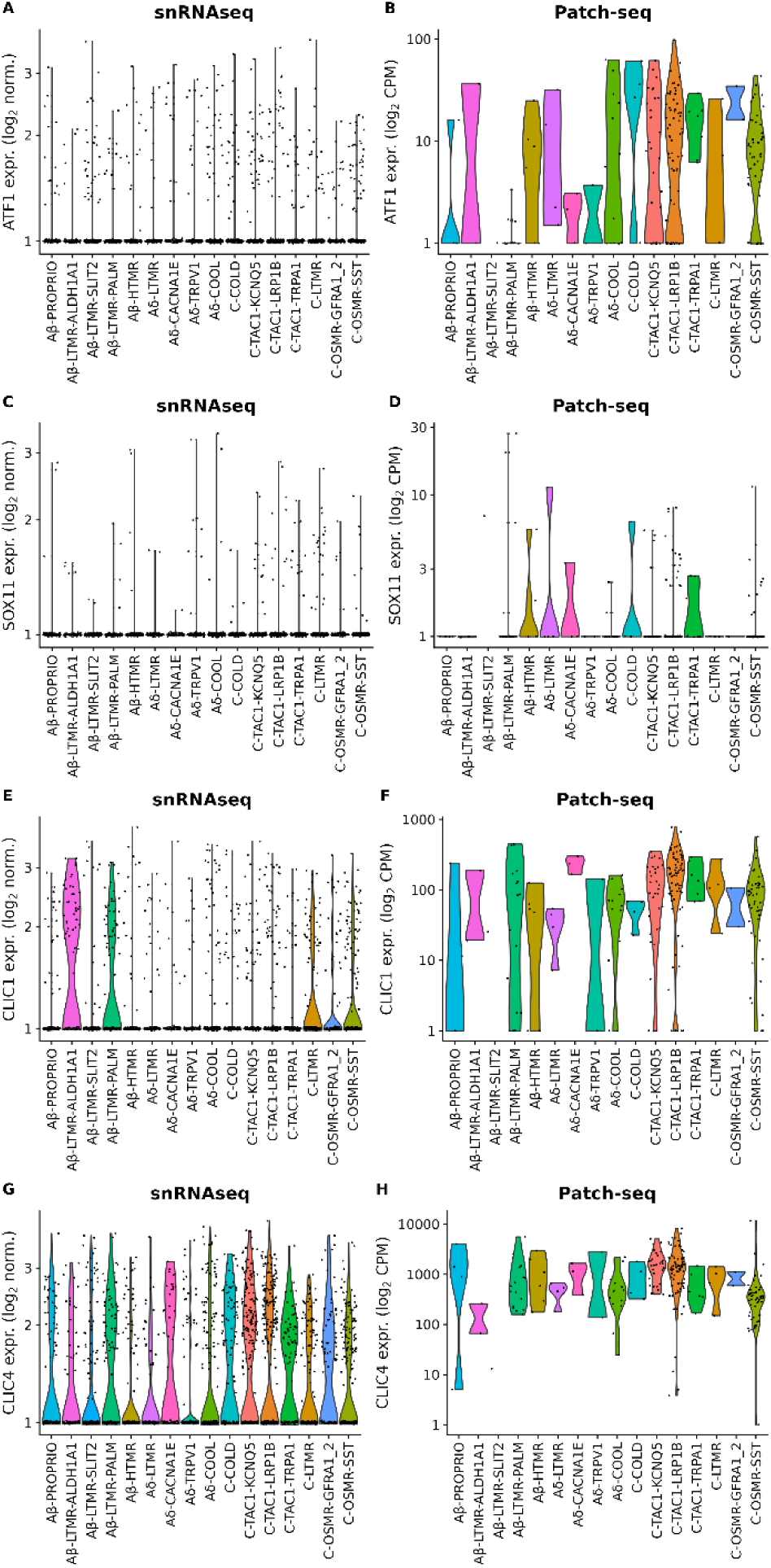
Comparison of injury-induced and culturing-related gene expression between snRNAseq and Patch-seq datasets. (A-D) Expression of known injury-induced genes across neuronal subtypes: ATF1 (A-B) and SOX11 (C-D) in snRNAseq (A, C) and Patch-seq (B, D) datasets. (E-H) Expression of culturing-associated genes: CLIC1 (E-F) and CLIC4 (G-H) in snRNAseq (E, G) and Patch-seq (F, H) datasets. Expression levels shown as log2-normalized counts (snRNAseq) or log2 CPM (Patch-seq). Each point represents a single cell/nucleus

**Figure S8:**
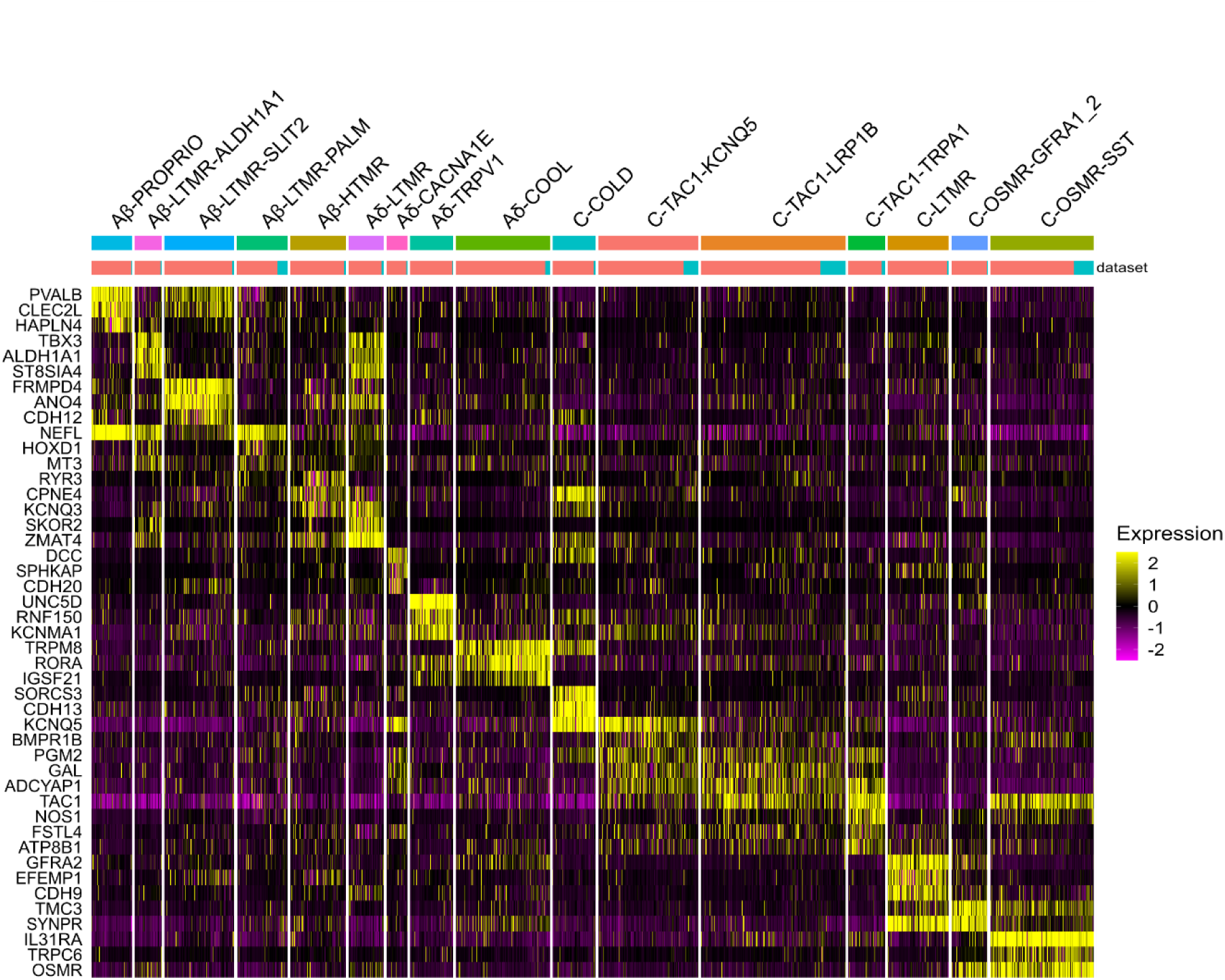
Expression of marker genes are conserved between Patch-seq and snRNAseq pig datasets. Heatmap showing the 3 most prominent gene products per neuronal identity. Upper bar indicates neuronal identity. Lower bar indicates dataset (snRNAseq, red; Patch-seq, blue).

**Figure S9:**
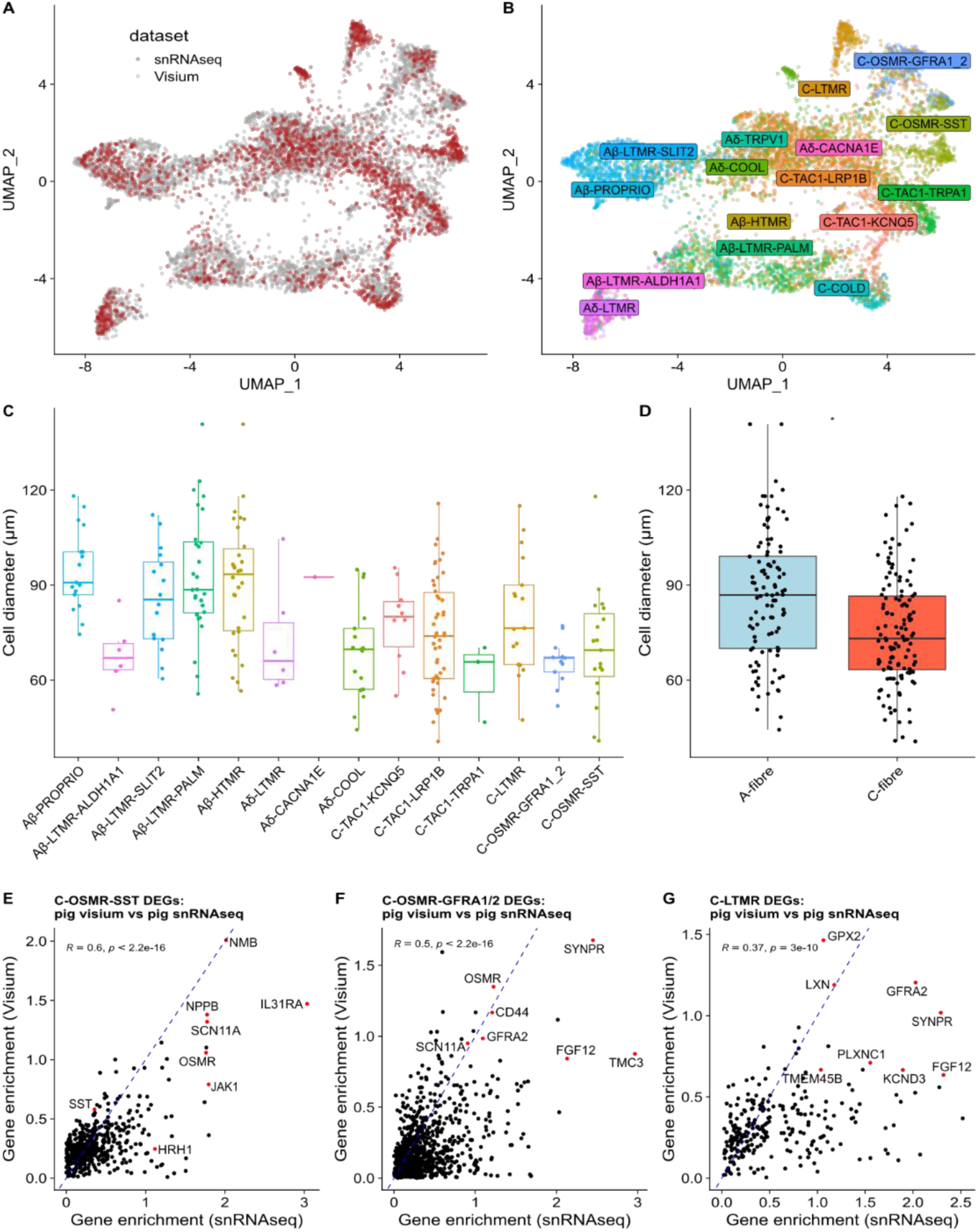
Spatial transcriptomics of pig DRGs corroborates cell type-enriched marker gene expression across technologies. **A**) UMAP representation of integrated gene expression from snRNAseq (red) and Visium-based spatial transcriptomics (grey). **B**) Same representation as in (A), but with colours distinguishing the mapped cell types. **C,D**) Cell diameters based on manual quantification of 230 DRG neurons from the Visium dataset illustrated as neuronal subtype (c), and at broad cell type resolution (85.5 ± 1.9 µm for A-fibers, 73.7 ± 1.6 µm for C-fibers, t=4.81, df=226.72, Welch’s t-test p-value = 2.72×10^-6^, mean ± SEM) (d). **E-G**) Illustration of concordance of cell type-enriched differential gene expression (DEGs) for C-OSMR-SST (E), C-OSMR-GFRA1/2 (F), and C-LTMR cells (G). Each dot reflects a gene, with x- and y-axis values indicating log2 fold changes of enrichment of target cell type (e.g., C-OSMR-SST cells) compared to all other cells in snRNAseq dataset (x-axis) and Visium spatial transcriptomics dataset (y-axis). Genes subset to those with positive enriched expression in both technologies. Inset values indicate Pearson correlations (R) and associated p-values. Genes denoted in red indicate cell type markers or otherwise notable genes.

**Figure S10:**
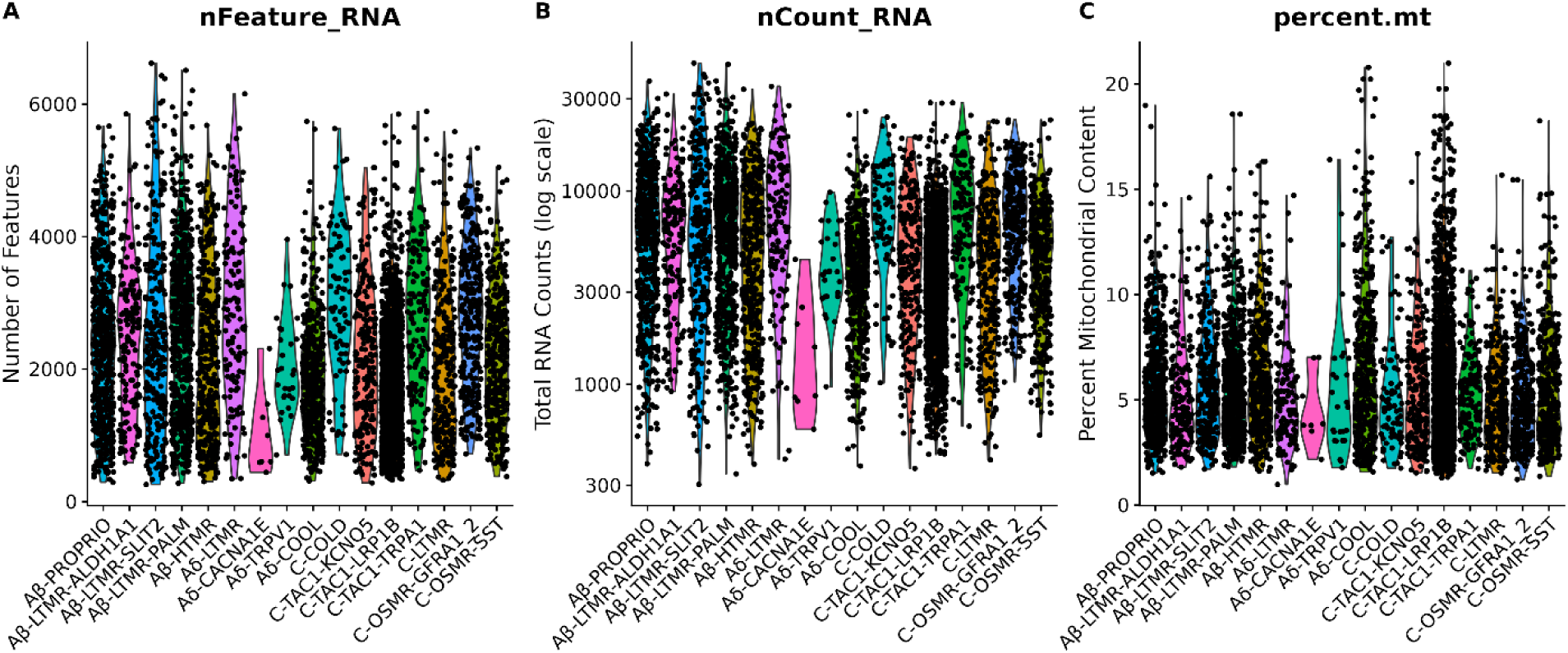
Sequencing metrics Visium data. **A-C**) Sequencing metrics for pig Visium dataset: a) number of reads per cell, b) number of genes per cell and c) fraction of mitochondrial reads per assigned neuronal identity. Each data point represents an individual cell.

**Figure S11:**
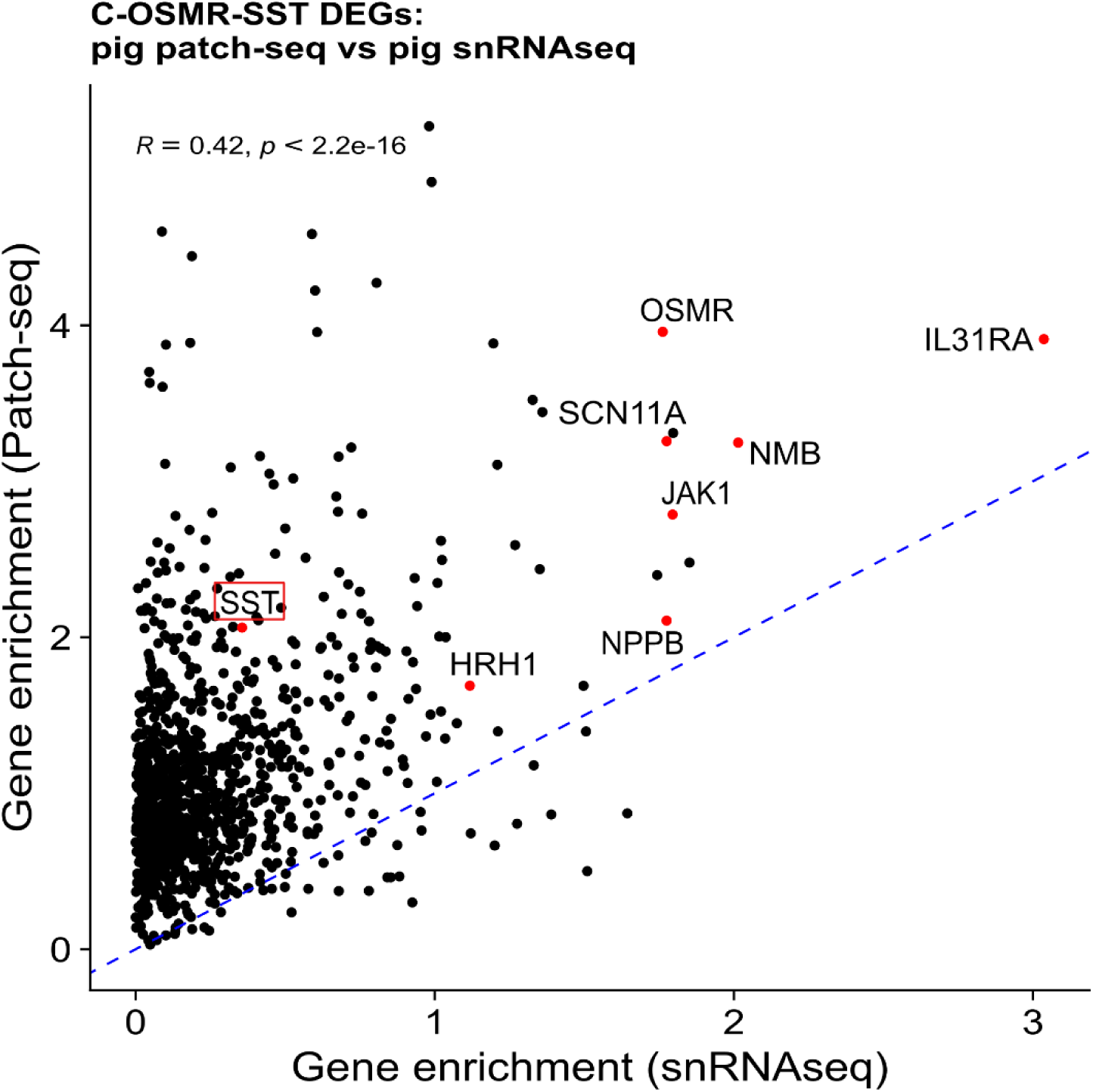
Concordance of differentially expressed genes in C-OSMR-SST across Patch-seq and snRNAseq. Concordance of differentially expressed genes (DEGs) in C-OSMR-SST cells across Patch-seq and snRNAseq datasets. Each point represents a gene, with the x-axis indicating gene enrichment log2 fold change in the snRNAseq dataset and the y-axis showing the same in the Patch-seq dataset, both derived from pig samples. Only genes with positive enrichment in both datasets are shown. Inset values indicate the Pearson correlation coefficient (R) and associated p-value. Genes denoted in red indicate cell type markers with SST highlighted despite relatively lower expression in the snRNAseq dataset.

**Figure S12:**
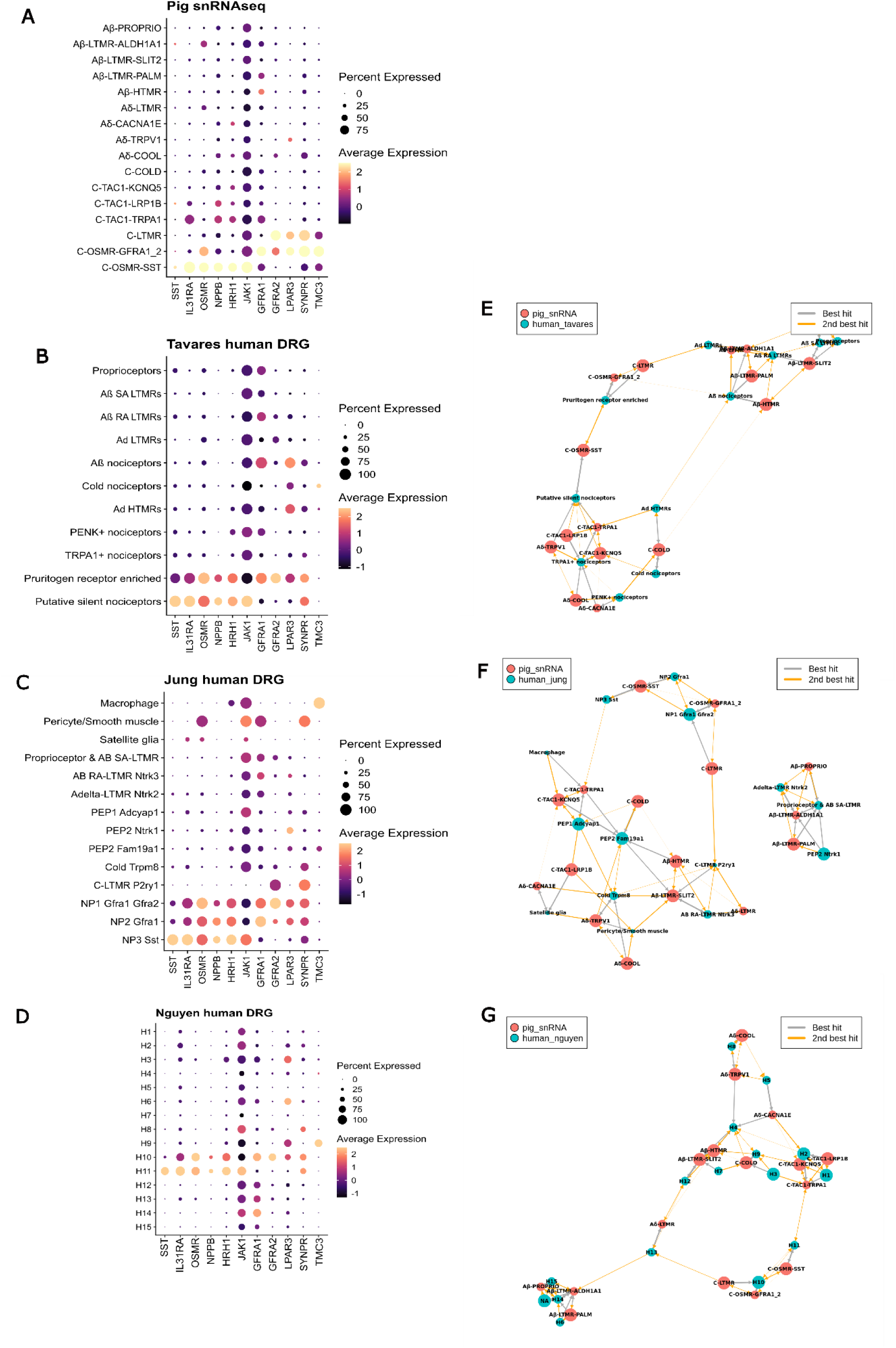
Pig and human DRG transcriptomic cell-type replicability. **A-D**) Dotplots showing expression of key marker genes for C-OSMR-SST neurons and related cell types across different datasets. The size of each dot represents the percentage of cells expressing the gene, while the color intensity indicates the average expression level. (**A**) Pig snRNAseq data collected in this study. (**B-D**) Human DRG data subsets from previously published cross-species atlas: (**B**) Tavares et al. human DRG dataset, (**C**) Jung et al. human DRG dataset, and (**D**) Nguyen et al. human DRG dataset. **E-G**) MetaNeighbor cluster graphs comparing pig snRNAseq cell types (red nodes) to human DRG cell types (blue nodes) from different studies in the cross-species atlas. (**E**) Comparison with Tavares et al. human DRG data. (**F**) Comparison with Jung et al. human DRG data. (**G**) Comparison with Nguyen et al. human DRG data.

**Figure S13:**
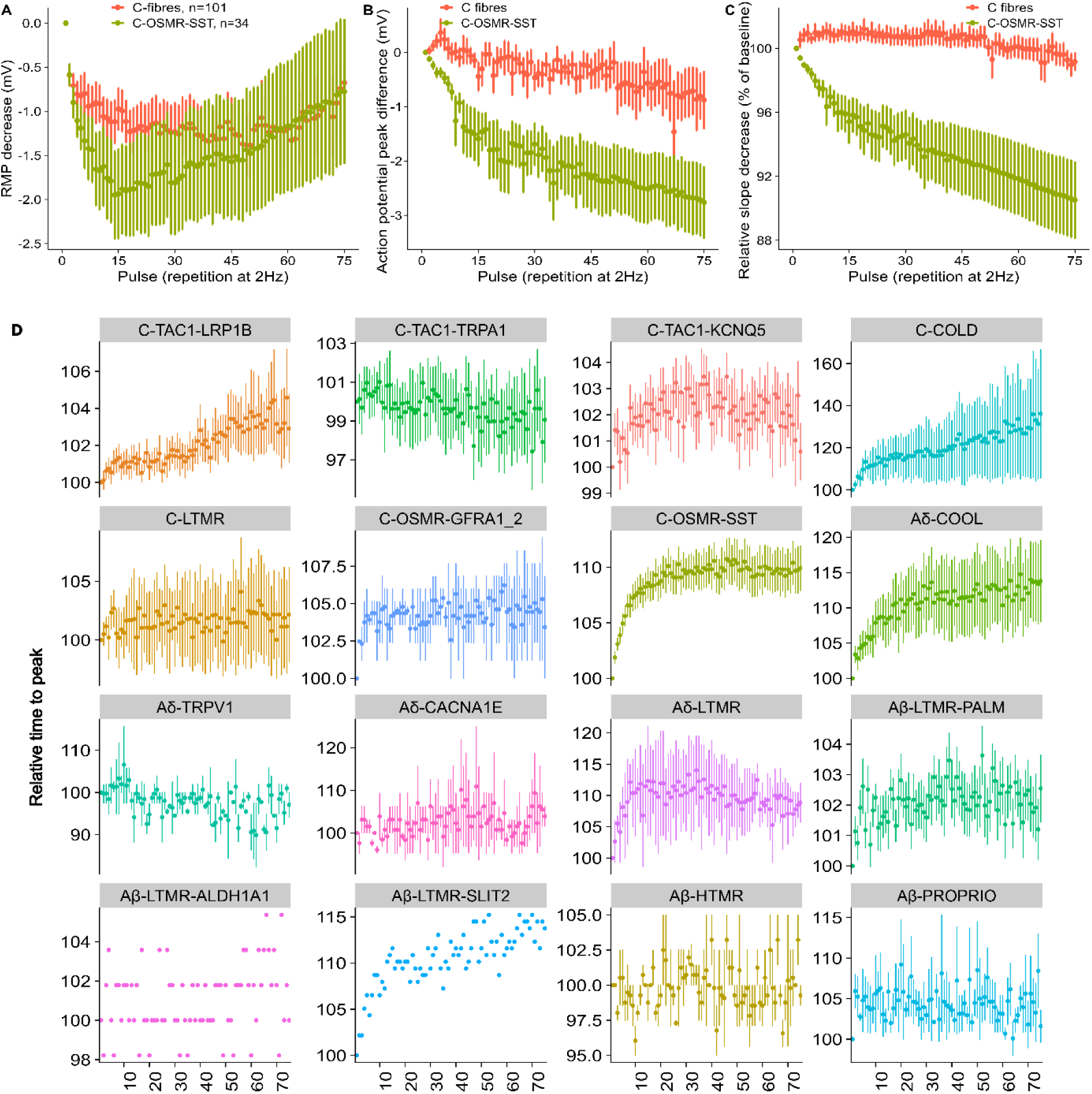
Extended electrophysiological characterization of CMi-fibers. Supplemental Data on functional biomarkers derived from *in vivo* microneurography experiments. **A**) Relative change in maximum action potential slope upon 75 rectangular 1000 pA stimulations with 10 ms and 2 Hz for C- OSMR-SST cells compared to all other Patch-seq characterized C-fiber neurons (mean change 0.906 ± 0.0235 SEM vs. 0.992 ± 0.00586 SEM; Mann-Whitney U = 2829.5, p < 1.733^-08^). **B**) Relative change in maximum action potential peak voltage upon 75 rectangular 1000 pA stimulations with 10 ms and 2 Hz for C-OSMR-SST cells compared to all other C-fiber neurons (mean change -3.08 ± 0.723 mV vs. -0.877 ± 0.540 mV; Mann-Whitney U = 2589.5, p < 9.855*10^-06^). **C**) Relative change in resting membrane potential upon 75 rectangular 1000 pA stimulations with 10 ms and 2 Hz for C-OSMR-SST cells compared to all other C-fiber neurons. **D**) Relative time to peak during 75 rectangular 1000 pA stimulations with 10 ms and 2 Hz for all cell types in the dataset. Error bars indicate SEM.

**Figure S14:**
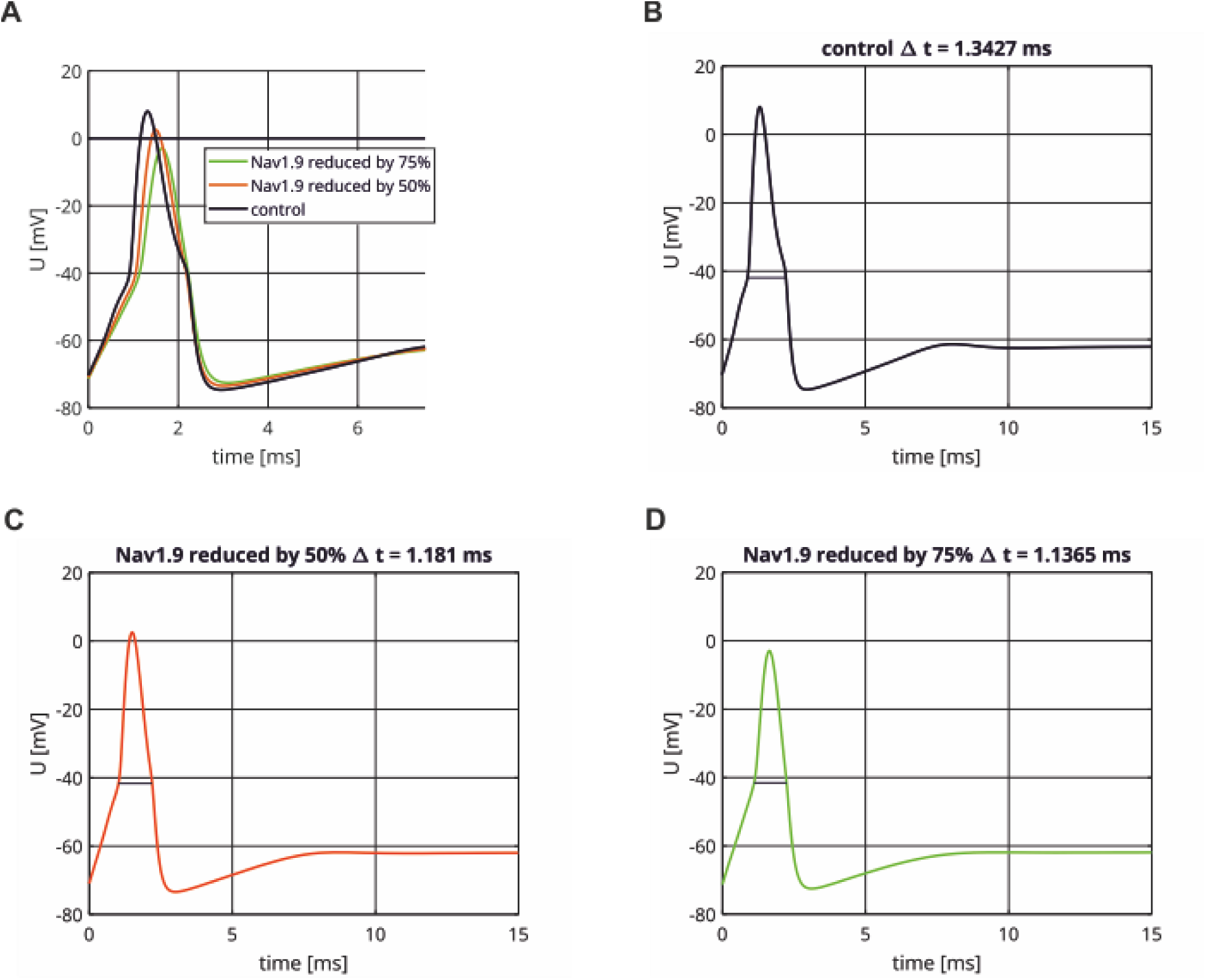
Simulated impact of SCN11A abundance on the AP morphology in C-OSMR-SST neurons. **(A):** Overlay of the three simulated AP with the relative abundance of SCN isoforms corresponding to their gene expression **B:** The black line (control) shows the simulated AP with the relative abundance of SCN isoforms corresponding to their gene expression. The maximal conductance attributed to SCN11A is stepwisely reduced to 50% (**C**) (red line) and 75% (green line) (**D**) compared to the control. The maximal conductances attributed to the other isoforms are increased proportionally to their expression, such that the total maximal sodium conductance remains unchanged. At the beginning of the simulation the neurons are in their resting state. The AP width (Δt) is measured at the threshold voltage and indicted by a horizontal line. Current injection starts at time 0 and lasts until the end of the simulation period. Maximal slopes: **A:** 213.2 mV/ms, **B**: 152.3 mV/ms. **C**: 109.6 mV/ms.

**Figure S15:**
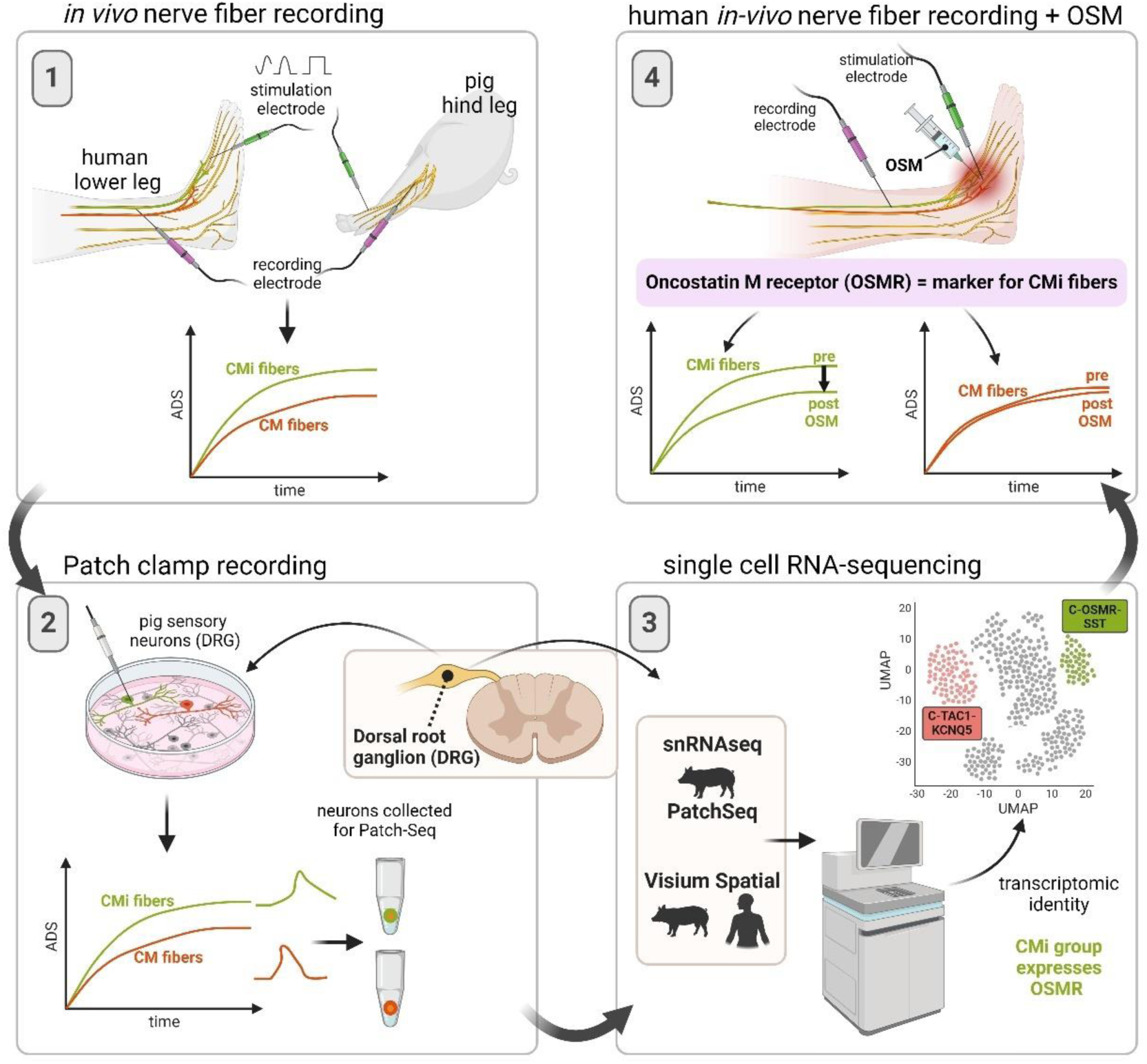
Graphical Abstract. Graphical abstract: Identifying the molecular identity of mechano-insensitive C-fibers. 1) Mechano-insensitive C- fibers (CMis) are traditionally defined functionally *in vivo* using specific electrical stimulation protocols applied to the skin. These protocols, activity dependent slowing (ADS), including sinusoidal and repetitive stimulation, distinguish CMis from mechano-sensitive C-fibers (CMs) in large mammals, including humans and pigs. 2) We used Patch-seq to characterize pig DRG neuron subtypes for whether they might be CMis. Current injection protocols applied *in vitro* were adapted from the stimulation protocols used to distinguish CMis *in vivo*. Following electrophysiological characterization, mRNA from neurons was harvested for sequencing. 3) We developed a novel multi-modal comparative/cross-species DRG neuronal taxonomy, encompassing single-nucleus RNAseq, Patch-seq, and Visium-based spatial transcriptomics in pigs and humans. This taxonomy enabled us to assess the putative CMi identity of each DRG molecular neuronal type. A distinct C-fiber molecular type, characterized by expression of *OSMR* and *IL31RA*, most likely reflects CMis. 4) Applying the OSMR ligand oncostatin M, to human subjects *in vivo* demonstrated a selective response to OSM by CMis but not CMs highlighting OSMR as key molecular marker of CMis.

**Supplementary Table 1.**
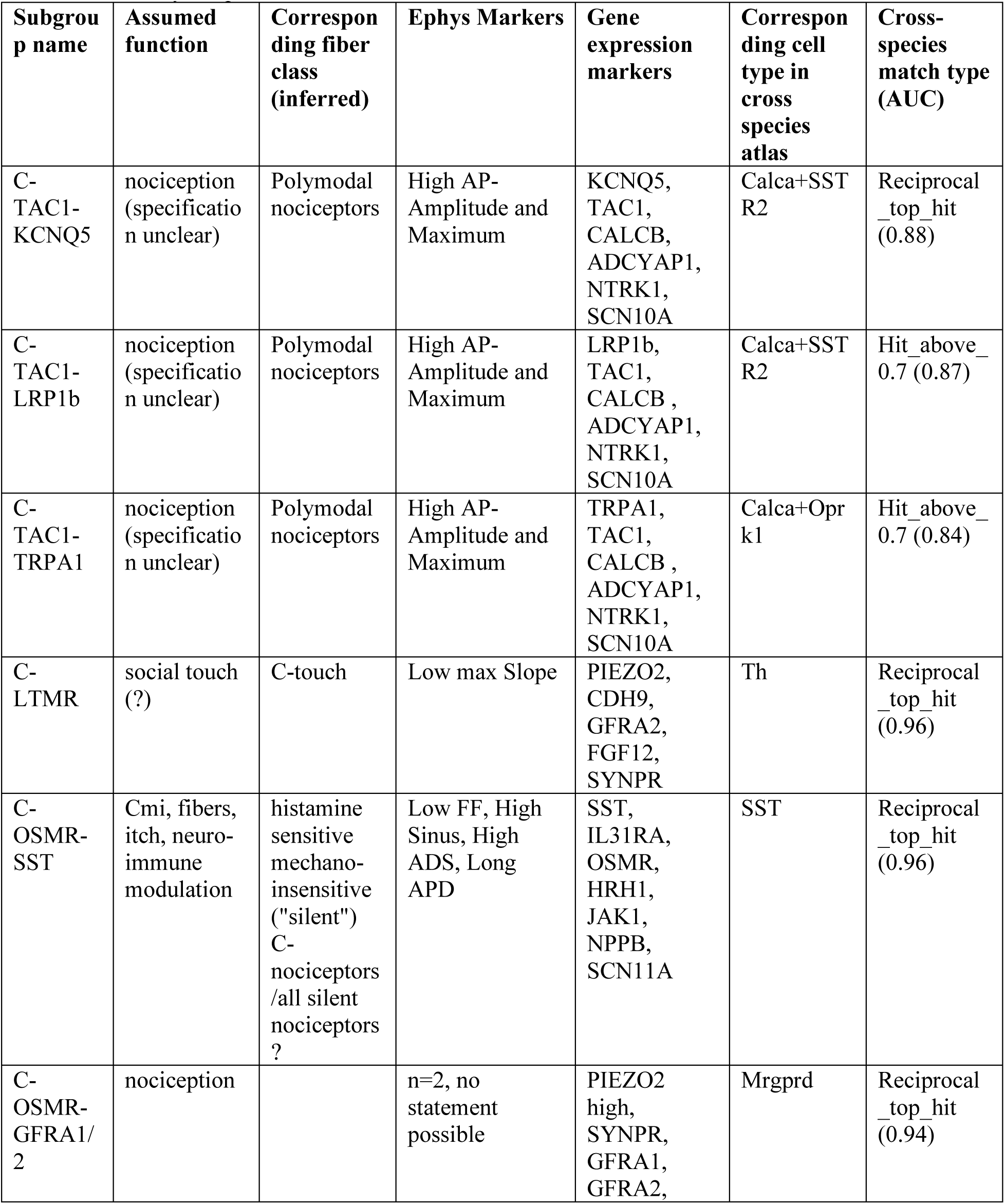
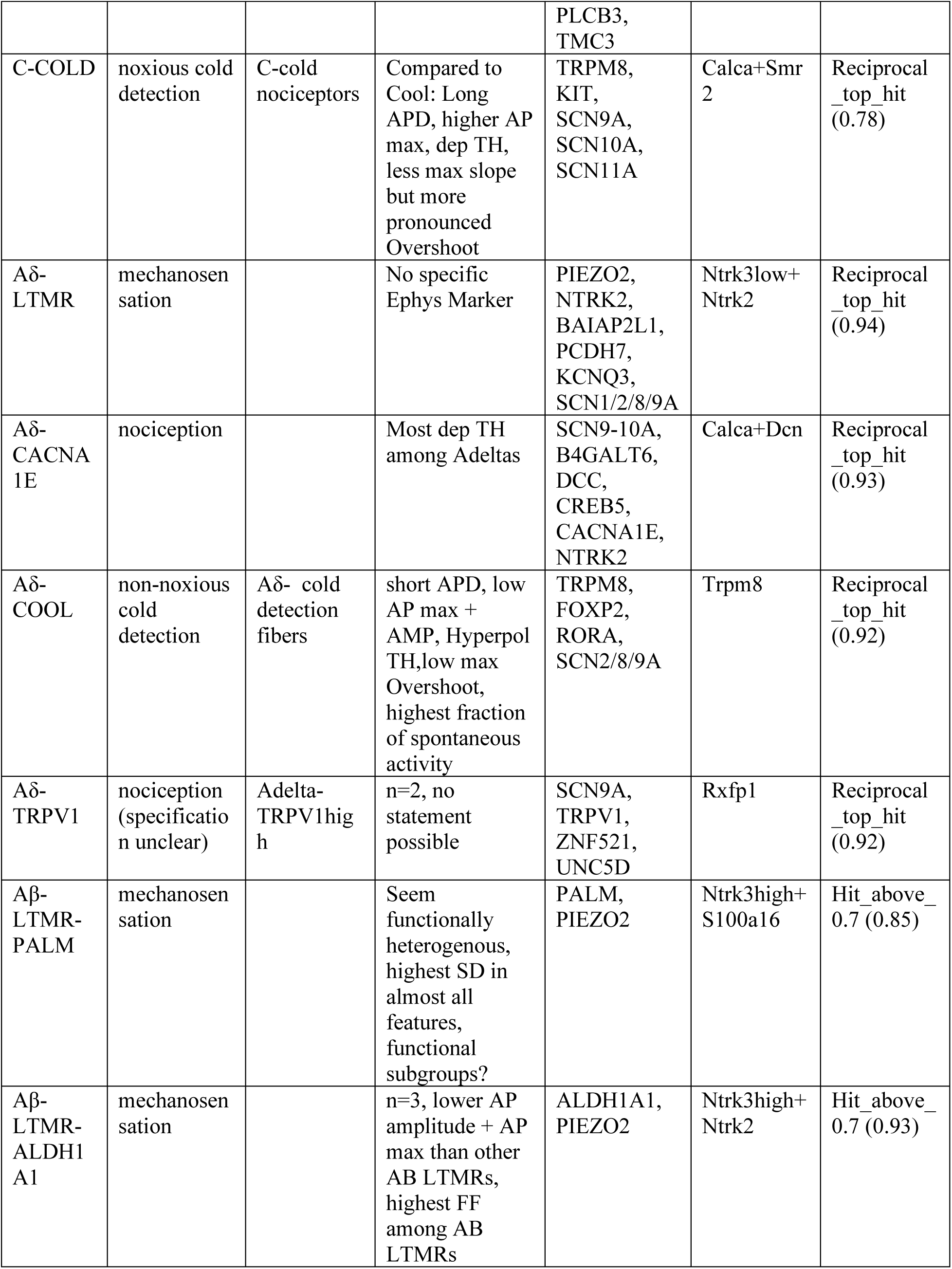
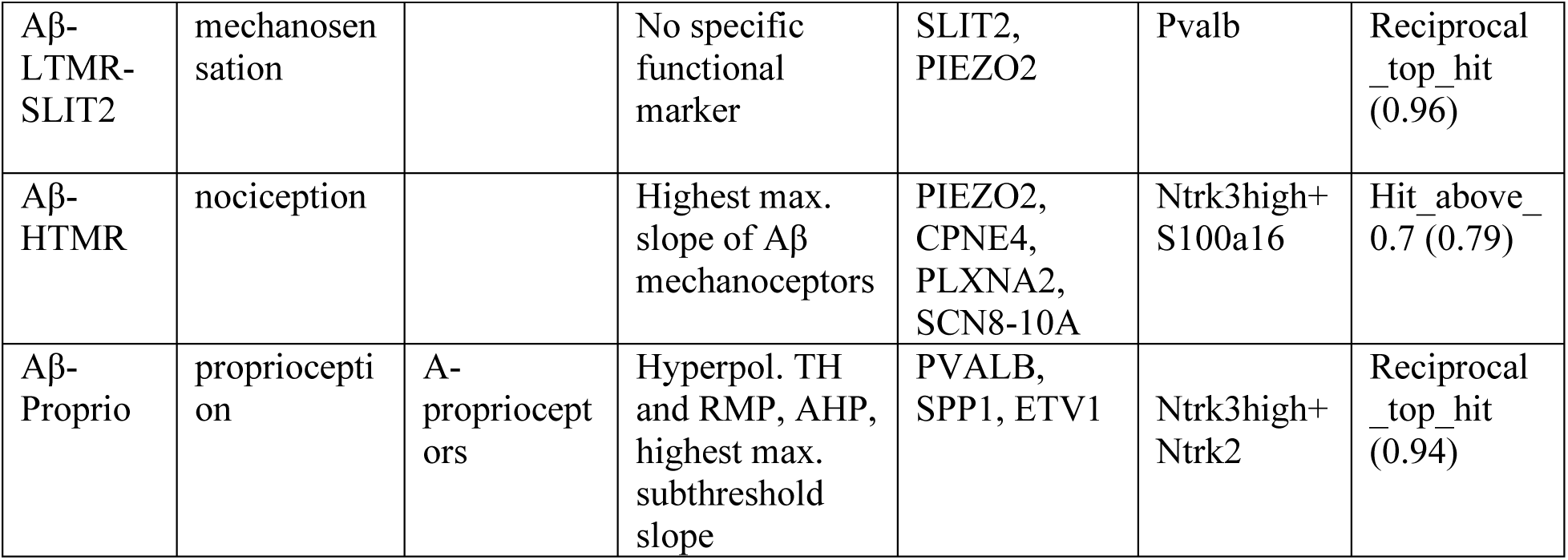
Overview on identified fiber classes with assumed function, corresponding fiber classes, electrophysiological markers, gene expression markers, and correspondence and MetaNeighbor-based match type and score to harmonized cross-species atlas. Reciprocal top hit indicates cell type pair is best matching cell type based on bi-directional taxonomy comparison.

